# Allosteric activation of the SPRTN protease by ubiquitin maintains genome stability

**DOI:** 10.1101/2024.11.26.625192

**Authors:** Sophie Dürauer, Hyun-Seo Kang, Christian Wiebeler, Yuka Machida, Dina S Schnapka, Denitsa Yaneva, Christian Renz, Maximilian J Götz, Pedro Weickert, Abigail C Major, Aldwin S Rahmanto, Sophie M Gutenthaler-Tietze, Lena J Daumann, Petra Beli, Helle D Ulrich, Michael Sattler, Yuichi J Machida, Nadine Schwierz, Julian Stingele

## Abstract

The DNA-dependent protease SPRTN maintains genome stability by degrading toxic DNA-protein crosslinks (DPCs). To understand how SPRTN’s promiscuous protease activity is confined to the cleavage of crosslinked proteins, we reconstitute the repair of DPCs including their modification with SUMO and ubiquitin chains, using recombinant human proteins. We discover that DPC ubiquitylation strongly activates SPRTN independently of SPRTN’s known ubiquitin-binding domains. Using protein structure prediction, MD simulations and NMR spectroscopy we reveal that ubiquitin binds to an interface at the back of SPRTN’s protease domain, promoting an active conformation. Replacing key interfacial residues prevents ubiquitin-dependent activation of SPRTN, which leads to genomic instability and cell cycle defects in cells expressing hypomorphic SPRTN variants that cause premature aging and liver cancer in Ruijs-Aalfs syndrome patients. Collectively, our results demonstrate that SPRTN activation is coupled to the modification of the crosslinked protein, explaining how specificity is achieved during DPC repair.

## INTRODUCTION

Cells invest in extensive repair mechanisms to ensure the fidelity of the genetic information stored in their DNA. Defective DNA repair results in mutagenesis and genome instability, major hallmarks of cancer, aging and aging-related diseases^1,2^. DNA repair pathways are regulated by sophisticated networks of post-translational modifications, deploying various enzymatic activities to target specific lesions in highly controlled manners^3,4^.

DNA-protein crosslinks (DPCs) are especially detrimental lesions that can arise enzymatically upon stabilization of covalent complexes between DNA-processing enzymes and their substrates^5^. For example, the transient covalent reaction intermediate formed between topoisomerase 1 and DNA can not only be stabilized by camptothecin and its clinical derivatives but also spontaneously by adjacent DNA damage^6,7^. Other enzymatic DPCs have important protective functions. The protein HMCES actively crosslinks to abasic (AP) sites within single-stranded DNA (ssDNA) to prevent AP site scission during DNA replication^8^. Additionally, various endogenous and environmental bifunctional reactive agents such as metabolic formaldehyde or ethanol-derived acetaldehyde inadvertently crosslink proteins to DNA^9,10^.

DPCs are highly toxic because large protein adducts block vital chromatin processes, including DNA replication and transcription^11–14^. The collision of the replication or transcription machinery with crosslinked proteins initiates repair^11–14^, which can additionally be triggered by global-genome mechanisms^10,15–18^. During the repair, the protein adduct is modified by ubiquitylation in all cases. During replication, ubiquitylation is carried out by the replisome-binding ubiquitin E3 TRAIP and the ssDNA-associated E3 RFWD3^19–21^, while the CRL4^CSA^ E3 ubiquitylates DPCs that have stalled elongating RNA polymerases^12–14^. During global-genome DPC repair, ubiquitylation is preceded by SUMOylation of the protein adduct^22^, which then leads to subsequent ubiquitylation by the SUMO-targeted ubiquitin E3s RNF4 and TOPORS^10,15–18^.

Following modification of the DPC^23^, repair proceeds by proteolytic destruction of the protein adduct. While the proteasome can degrade DPCs^10,15,16,19,20^, they are also targeted by specialized DPC proteases^24,25^. SPRTN, the major human DPC protease, targets DPCs during replication and in global-genome pathways^10,19,26,27^. Loss of SPRTN is lethal in virtually all human cell lines^28^ and leads to dramatic genome instability and early embryonic lethality in mice^29^, highlighting SPRTN’s critical role in repairing endogenously arising DNA damage.

SPRTN features a metalloprotease domain at the *N*-terminus, which, together with the ssDNA-binding zinc-binding domain (ZBD), forms the conserved SprT domain (Figure 1A)^30,31^. SPRTN’s protease activity is not specific to certain amino acid sequences, which is perfectly suited to target diverse types of protein adducts, but also requires tight regulation to prevent accidental cleavage of chromatin proteins. The SprT domain is followed by a basic region (BR) that specifically interacts with double-stranded DNA (dsDNA)^26^. ZBD and BR couple SPRTN activity to the recognition of DNA structures that contain both single- and double-stranded features, including ssDNA-dsDNA junctions^32^. Such junctions arise when replicative polymerases stall at DPCs during replication^19^. Despite these insights, the recognition of ssDNA-dsDNA junctions or similar structures cannot explain how specificity is achieved during SPRTN activation, given that these DNA structures are not specific to sites of DPC formation. On the contrary, they are common throughout the genome, for example on the lagging strand during DNA replication. Therefore, it remains unknown how SPRTN’s protease activity is confined to cleavage of crosslinked proteins. Presumably, SPRTN activation must involve a conformational change^26^, given that the only available structure of the SprT domain revealed a closed conformation, in which the ZBD domain restricts access to SPRTN’s active site^31^.

**Figure 1.**
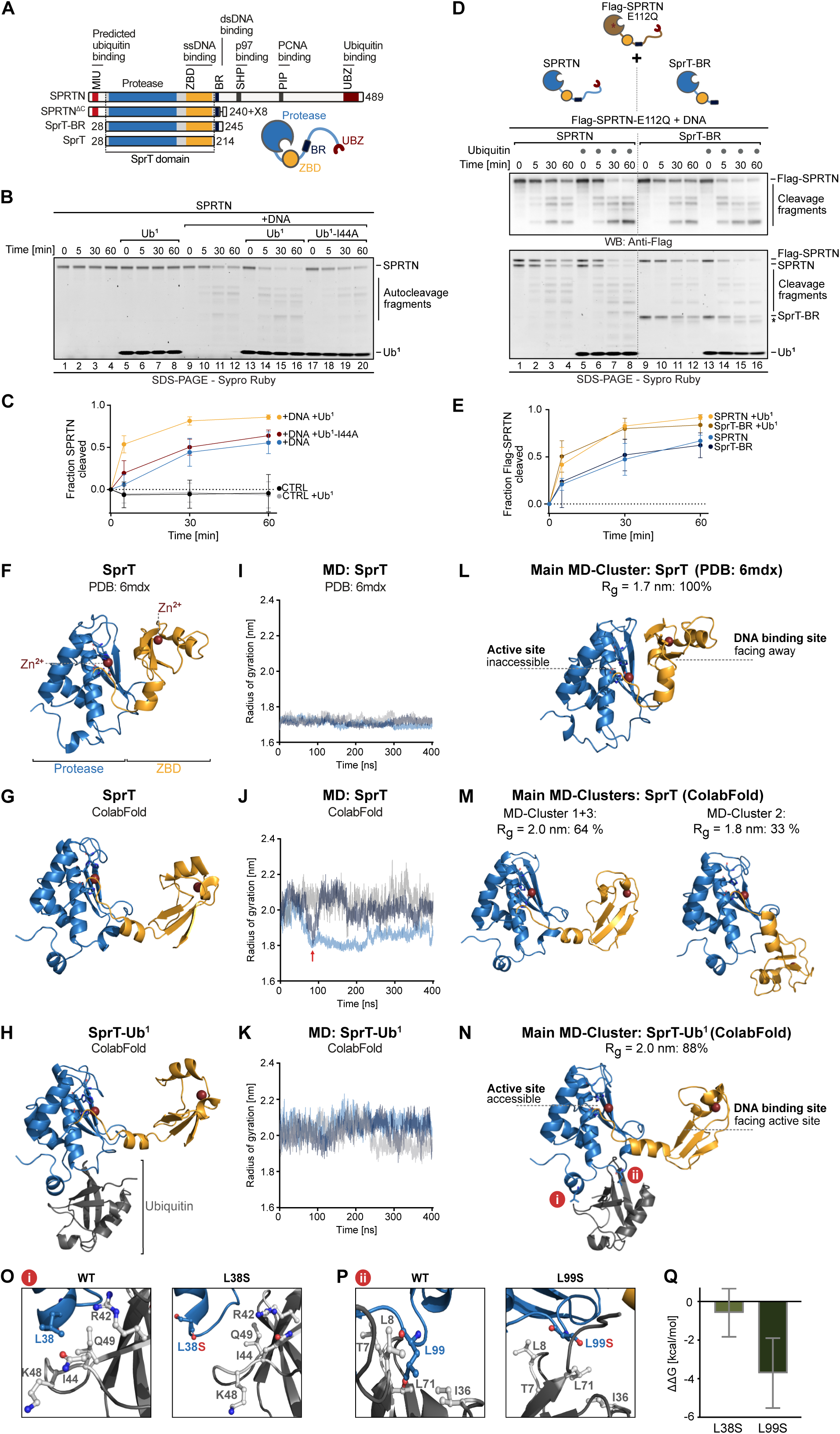
Ubiquitin promotes an open SPRTN conformation. (A) Schematic of SPRTN’s domain structure and truncated variants, featuring motif interacting with ubiquitin (MIU), protease domain, zinc-binding domain (ZBD), basic region (BR), SHP box for p97-binding, PCNA-interacting motif (PIP) and ubiquitin-binding zinc finger (UBZ). SPRTN^ΔC^ is caused by a frameshift mutation resulting in a variant composed of SPRTN’s N-terminal 240 residues followed by eight additional amino acids (X8). (B) SPRTN (250 nM) was incubated alone or in the presence of activating Virion DNA (11.14 nM) with or without mono-ubiquitin (Ub^1^ or Ub^1^-I44A, 10 µM). (C) Quantification of SPRTN autocleavage assay shown in (B), depicting the mean ± SD of five independent experiments. (D) SPRTN or SprT-BR (250 nM) was incubated in the presence of Virion DNA (11.14 nM) and Flag-SPRTN-E112Q (250 nM) with or without Ub^1^ (10 µM). SprT-BR autocleavage fragment is marked with an asterisk. (E) Quantification of SPRTN autocleavage assay shown in (D), depicting cleavage of Flag-SPRTN-E112Q as the mean ± SD of three independent experiments. (F-H) Experimental structure of SPRTN’s SprT domain (SPRTN^aa28–214^, PDB: 6mdx) (F), ColabFold predicted structure of SprT (G) and ColabFold predicted structure of a SprT-Ub^1^ complex (H). (I-K) Radius of gyration (Rg) of the indicated structures over 400 ns of molecular dynamics (MD) simulation. Each curve represents an independent MD trajectory (n=3). (L-N) Main MD-clusters of the indicated structures during MD simulation for 400 ns, generated from three independent trajectories. For SprT (ColabFold predicted) two of three main MD-clusters are depicted. Rg correlating frequencies among all performed simulations are labeled above structures. (O-P) Zoom-In to regions **i** and **ii** of the SprT-Ub^1^ complex (N), showing amino acids of ubiquitin (in grey) surrounding residue Leu38 (O) or L99 (P) of SPRTN (in blue) in the wild-type (WT) protein (left) and upon L38S or L99S replacement, respectively (right). (Q) SprT+Ub^1^ binding energy difference (ΔΔG) between SprT-L38S or -L99S and WT protein obtained from alanine scanning. Bar graphs show the mean ± SD of 301 snapshots from PBSA calculations for the central structure of the largest cluster. See also Figure S1.

In addition to its DNA-binding domains, SPRTN bears interaction motifs for binding to the segregase p97 (SHP box) and PCNA (PIP box)^33–36^ but neither is required for SPRTN’s DPC repair function^10,19,29^. Furthermore, SPRTN carries a *C*-terminal ubiquitin-binding zinc finger (UBZ), promoting mono-ubiquitylation of SPRTN, which has been linked to its inactivation^37^. A motif interacting with ubiquitin (MIU) has been predicted at SPRTN’s very *N*-terminus but has not been experimentally confirmed yet^38^. The precise role of ubiquitin in SPRTN-mediated DPC cleavage has remained controversial. Cleavage of a model DPC by SPRTN can be observed in frog egg extracts even if the protein adduct has been treated with formaldehyde, to prevent efficient ubiquitylation^19^. Nonetheless, ubiquitylated DPCs accumulate upon SPRTN depletion^39^, suggesting that they are substrates of the protease. It has been speculated that the UBZ may help to recruit SPRTN to ubiquitylated DPCs. In agreement, the UBZ domain supports efficient DPC cleavage in frog egg extracts and also in cells^10,19^. Surprisingly however, the UBZ domain is not essential for SPRTN’s function. Patients with Ruijs-Aalfs syndrome (RJALS) express truncated versions of SPRTN that lack the *C*-terminal part of the enzymes including the UBZ (SPRTN^ΔC^; Figure 1A), either alone or in combination with active site mutations^38^. RJALS patients suffer from premature aging and predisposition to liver cancer^38^, phenotypes that are recapitulated in a mouse model of reduced SPRTN function^29^. Yet, the truncated SPRTN patient variants are clearly compatible with life, in contrast to full loss of SPRTN. In agreement, the severe growth defects associated with SPRTN loss in conditional mouse knock-out cells are fully rescued by expression of truncated SPRTN^40^. Hence, it has remained enigmatic how SPRTN patient variants target DPCs in the absence of the UBZ and, more generally, whether and how SPRTN activity is regulated by DPC ubiquitylation.

Here, we investigate the role of ubiquitin in SPRTN activation using a multi-pronged approach, encompassing the biochemical reconstitution of DPC modification and repair, molecular dynamics (MD) simulations, NMR experiments and cellular assays. We find that ubiquitin, particularly DPC ubiquitylation, strongly activates SPRTN. The ubiquitin-dependent activation of SPRTN occurs independently of its UBZ, but instead requires a novel ubiquitin-binding interface at the back of its protease domain. This interface is critically required in cells expressing *C*-terminally truncated RJALS patient variants to maintain cellular fitness and to prevent chromatin bridges, micronuclei and cell cycle defects. Collectively, our results reveal a novel regulatory principle that confines SPRTN’s protease activity by linking its activation to the ubiquitylation of the DPC and explain how residual SPRTN function is maintained in RJALS patients.

## RESULTS

### Ubiquitin stimulates SPRTN’s DNA-dependent protease activity by promoting an open conformation

To explore the relationship between SPRTN and ubiquitin, we monitored DNA-dependent autocleavage of recombinant human SPRTN (Figure S1A) in the presence and absence of ubiquitin; in this type of assay, autocleavage occurs *in trans*^26^. Upon addition of ubiquitin, SPRTN autocleavage was enhanced in the presence of DNA (Figure 1B, compare lanes 9-12 and 13-16, Figure 1C for quantification). In the absence of DNA, no autocleavage was observed independent of whether ubiquitin was present or not (Figure 1B, lanes 1-8, Figure 1C for quantification), suggesting that ubiquitin promotes SPRTN’s DNA-dependent protease activity. The stimulating effect required ubiquitin’s Ile44 residue (Figure 1B, compare lanes 13-16 and 17-20, Figure 1C for quantification). Given that most ubiquitin-binding domains (UBDs) interact with ubiquitin by binding to the hydrophobic patch around Ile44^41,42^, this observation indicated to us that a UBD in SPRTN is mediating the stimulating effect of ubiquitin. Notably however, deletion of SPRTN’s UBZ or its predicted MIU domain had no effect on the ubiquitin-dependent activation (Figure S1B, Figure S1C for quantification). Even a minimal catalytically active SPRTN fragment (SprT-BR, aa28-245) displayed further activation upon addition of ubiquitin (Figure S1D, Figure S1E for quantification). Because autocleavage fragments produced by truncated SprT-BR are difficult to compare with fragments originating from full-length SPRTN, we additionally assessed *in trans* cleavage of a catalytically inactive Flag-tagged SPRTN (Flag-SPRTN-E112Q). SPRTN and SprT-BR cleaved Flag-SPRTN-E112Q with comparable efficiency in the absence of ubiquitin and were further activated to the same extent upon addition of ubiquitin Figure 1D, top panel, compare lanes 1-4 with lanes 9-12, and lanes 5-8 with lanes 13-16, Figure 1E for quantification). The stimulating effect of ubiquitin on truncated SprT-BR suggested the presence of an additional ubiquitin-binding site within this region of SPRTN (Figure 1A).

To explore this possibility, we used ColabFold^43^ to predict complexes between SprT-BR and ubiquitin. In the top-ranked model (Rank_1), ubiquitin was predicted to interact via its Ile44 patch with an interface at the back of the SprT domain (Figure S1F-G), hereafter referred to as ubiquitin-binding interface at the SprT domain (USD). Interestingly, the SprT domain was predicted in all models to adopt an open conformation with a highly accessible active site facing the ZBD’s DNA binding site. A similar conformation was predicted in the absence of ubiquitin, in stark contrast to the published crystal structure of the SprT domain (PDB:6mdx^31^) that shows a closed conformation with the ZBD restricting access to the active site (Figure 1F-H). Next, we set out to gain insights into whether the predicted open conformation of the SprT domain may be in equilibrium with the closed conformation and thereby enable interaction with ubiquitin. Therefore, we conducted all-atoms MD simulations using the crystal structure or ColabFold-based predictions of the SprT domain, alone or in combination with ubiquitin, as starting points (Figure 1I-K and Figure S1H). The compact conformation observed in the crystal structure remained largely unchanged over the entire 400 ns timeframe in three independent simulations (Figure 1I). To reveal the predominant SprT conformations within the different simulations, we employed RMSD-based clustering (Figure 1L-N). Simulations of the crystal structure consisted of a single cluster with a closed conformation (radius of gyration (Rg)=1.7 nm, Figure 1L). In contrast, simulations of the ColabFold-predicted SprT structure revealed larger conformational changes during the simulations (Figure 1J). We observed transient collapses of the SprT domain to a compact conformation with low Rg (Figure 1J, red arrow). The collapse was either followed by rapid reopening of the structure (Figure 1J, dark blue trace) or prolonged collapse lasting several hundred nanoseconds (Figure 1J, light blue trace). Indeed, clustering revealed three main clusters. The first and third cluster exhibited an open conformation (Rg=2.0 nm, Figure 1M, left), whereas the second cluster showed a closed conformation (Rg=1.8 nm, Figure 1M, right). The presence of ubiquitin prevented conformational transitions to the closed structure within the SprT domain (Figure 1K) and the simulations predominantly remained in an open conformation (Rg=2.0 nm, Figure 1N). Notably, ubiquitin binding to the USD interface of the SprT remained stable across all three independent simulations (Figure 1K). These data indicated to us that ubiquitin binding at the SprT domain may promote SPRTN activation by stabilizing an open conformation of the enzyme with an accessible active site.

Next, we wanted to determine amino acid residues within the USD interface that are important for ubiquitin-binding. In the predicted SprT-ubiquitin complex, Leu38 and Leu99 of SPRTN appeared to mediate the interaction via hydrophobic interactions involving multiple amino acids within ubiquitin’s hydrophobic Ile44- and Ile36-patch, respectively (Figure 1O-P and Figure S1I). To assess the effect of replacing either leucine residue with a hydrophilic serine (L38S, L99S), we conducted free energy end-point calculations using MMPBSA in conjunction with alanine scanning (see Methods for details), which enabled us to quantify the effect of each leucine-to-serine replacement to the overall binding affinity of the SprT-ubiquitin complex. We calculated a decrease in binding affinity of around 0.6 kcal/mol for the L38S replacement and 3.74 kcal/mol for L99S (Figure 1Q). This effect is explained by replacement of Leu38 and Leu99 resulting in the loss of hydrophobic contacts to ubiquitin’s Ile44 and Ile36 patch, respectively (Figure 1O-P and Figure S1I).

Collectively, our results suggest that ubiquitin may stimulate SPRTN activity by interacting with the USD interface of SPRTN’s SprT domain, potentially stabilizing an open – presumably active – conformation of the protease.

To experimentally test whether ubiquitin binds to the USD interface and whether ubiquitin binding affects SPRTN’s interaction with DNA, we used NMR spectroscopy. Heteronuclear single quantum coherence (HSQC) spectra of a catalytically inactive SprT-BR construct (E112Q) show well-dispersed peaks, indicative of the presence of folded regions (Figure S1J and S2A). The crowded middle region corresponds to the unstructured linkers and the *C*-terminal BR region. Chemical shift perturbations (CSP) difference seen in spectral overlays of SprT-BR with a construct lacking the protease domain (ZBD-BR, Figure S2A) highlight some differences in the linker between protease and ZBD (aa151-160) and on the β-sheet fold of the ZBD (Figure S2B). Nonetheless, we were able to transfer many chemical shifts from our previous analysis of the ZBD-BR construct^32^. The large CSPs of residues aa151-160, presumably reflect interaction of this region with the protease domain (Figure S2B). Due to the non-optimal sample stability of SprT-BR, we could not assign the individual resonances of the protease domain. However, by exclusion, the resonances belonging to the protease domain could readily be distinguished from those in the ZBD and BR (see Figure 2A for the Trp ε1 resonances).

**Figure 2.**
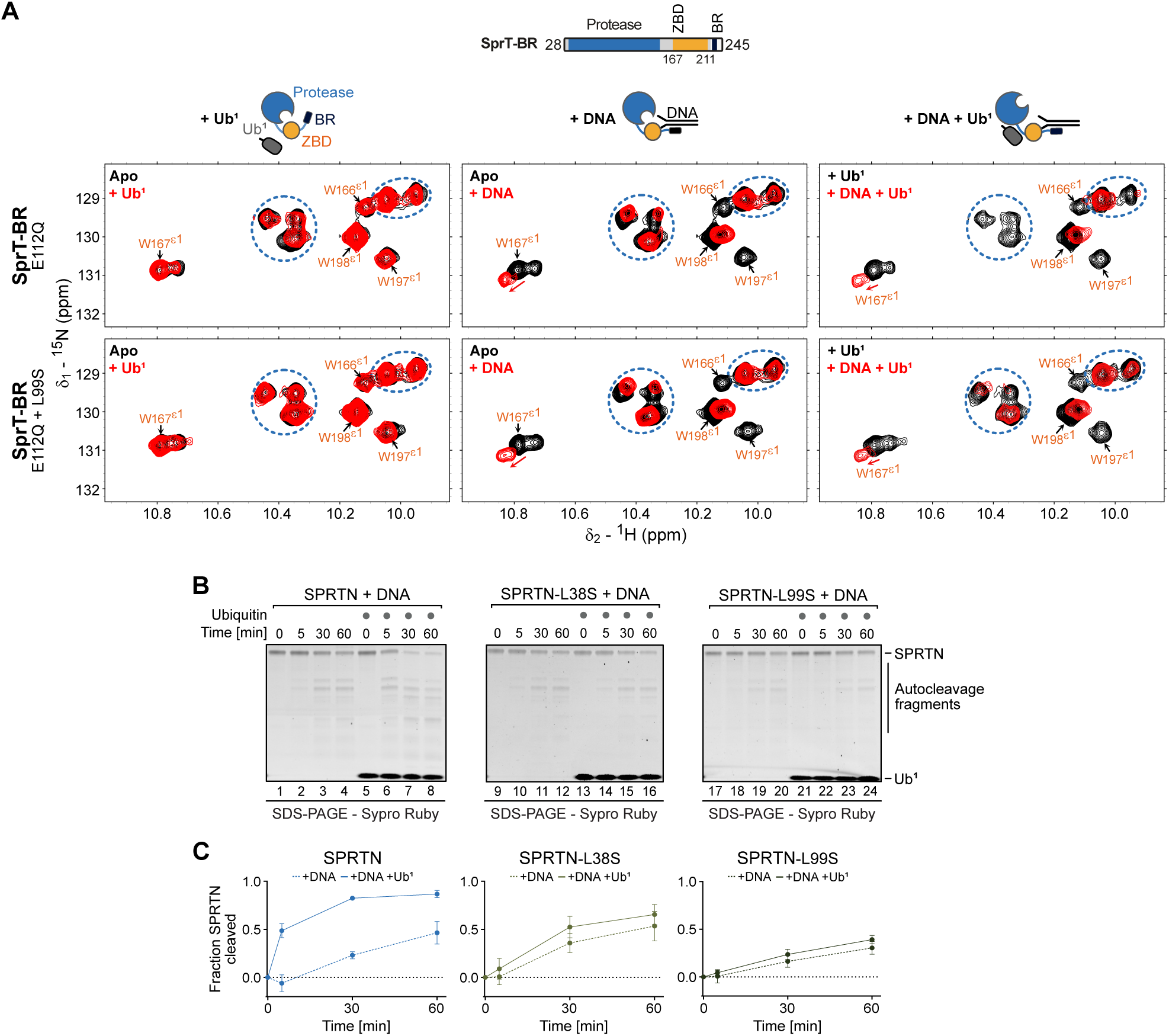
Ubiquitin binds to the SprT domain via the USD. (A) Comparison of NMR spectral (Trp ε1 amide signals in ^1^H,^15^N-HSQC experiments) of catalytically inactive SprT-BR-E112Q (upper row) and SprT-BR-E112Q+L99S (lower row) alone (black) (=Apo), with mono-ubiquitin (Ub^1^) (5x molar excess) (red) (left), with dsDNA (2x molar excess) (red) (middle), and Ub^1^-bound (5x molar excess) SprT-BR-E112Q and SprT-BR-E112Q+L99S alone (black) and in presence of dsDNA (2x molar excess) (red) (right), corresponding to cartoons above. Resonances corresponding to the Trp ε1’s in the zinc-binding domain (ZBD) are labeled (orange) or shown as asterisk when broadened. Trp ε1’s in the protease domain are highlighted with blue circles. Full spectra are shown in Figure S2. (B) Indicated variants of SPRTN (250 nM) were incubated in the presence of Virion DNA (11.14 nM) with or without Ub^1^ (10 µM). (C) Quantification of SPRTN autocleavage assay shown in (B), depicting the mean ± SD of three independent experiments. See also Figure S2 and Figure S3.

To examine the effects of ubiquitin and DNA binding on SPRTN’s conformation, we focused our analyses on Trp ε1 ^1^H,^15^N resonances, which we could unambiguously assign to the ZBD (Figure 2A, assigned, orange labels, and S2C) and protease domain (Figure 2A, blue circles, and S2C), respectively. To assess contributions of the USD interface, we determined the effect of replacing Leu99 with serine. NMR spectra of SprT-BR and the corresponding L99S construct superimposed very well (Figure S3A), except for those resonances in vicinity to the mutation site, indicating structural integrity is not affected upon replacement of Leu99. Additionally, Trp ε1 amide resonances were not affected by the mutation. Upon adding ubiquitin in five-fold excess, we observed changes in the protease domain of SprT-BR spectra (Figure S2C, blue boxes, top panels). Trp ε1 resonances located in the protease domain were slightly broadened (Figure 2A, top left panel, blue circles). In the L99S variant, the effects of ubiquitin addition were significantly reduced, implying that they correspond to ubiquitin binding to SPRTN’s USD interface (Figure 2A, bottom left panel, blue circles). While the ubiquitin-induced effects were subtle and mostly affected resonances corresponding to the protease domain, we also observed line-broadening for signals corresponding to ZBD (Figure S3B, left). These residues are again located in the β-sheet close to the linker region (Figure S3B, right panel). On the other hand, the addition of an activating DNA structure in two-fold excess led to major spectral changes in ZBD-BR regions, showing significant line-broadening, which was comparable between wild type (WT) and L99S constructs for the Trp ε1 resonances (Figure 2A, middle panels). This was also clearly reflected in the full spectral comparison (Figure S2D, middle panels). These results demonstrate that alteration of the USD does not affect DNA binding.

Strikingly, upon combined addition of both DNA and ubiquitin, severe line-broadening of the Trp ε1 resonances was observed in SprT-BR that was more pronounced than the individual effects of ubiquitin or DNA binding (Figure 2A, top, right panel). The effect was also clearly noticeable in the full spectra (Figure S2C, bottom, left panel), suggesting that the simultaneous binding of ubiquitin and DNA has synergistic effects on SPRTN’s conformation. These effects were virtually absent in the L99S variant for the Trp ε1 resonances (Figure 2A, bottom, right panel) and, overall, in the full spectrum (Figure S2C, bottom, right panel). Consistently, addition of ubiquitin with a mutated Ile44 patch had little effect (Figure S3C-D). Collectively, our NMR data indicate that ubiquitin amplifies the effects of DNA binding on SPRTN conformation allosterically by binding to the USD interface at the back of the protease domain.

To directly test whether a lack of ubiquitin-binding and the associated conformational changes within SPRTN translate to differences in enzymatic activity, we produced full-length SPRTN proteins with an L38S or L99S substitution (Figure S1A). While both variants showed DNA-induced autocleavage activity, they were either only mildly (L38S) or not at all (L99S) activated by ubiquitin, in contrast to the WT enzyme (Figure 2B and Figure 2C for quantification).

Collectively, these results demonstrate that ubiquitin activates SPRTN by binding to the USD interface. Our MD simulations and NMR results further suggest a model wherein ubiquitin binding to the USD stabilizes an open conformation of the enzyme in the presence of DNA.

### Ubiquitin chains are particularly potent at SPRTN activation

To gain deeper insight into the ubiquitin-dependent activation of SPRTN, we explored whether ubiquitin chains differ from mono-ubiquitin in their ability to activate. Therefore, we compared activation of SPRTN by mono-ubiquitin (Ub^1^) with linear (M1-), K48- and K63-linked tetra-ubiquitin chains (Ub^4^) in SPRTN autocleavage assays. All tetra-ubiquitin chains stimulated SPRTN activity to a substantially stronger degree than mono-ubiquitin (Figure 3A and Figure 3B for quantification, chain concentrations refer to the concentration of mono-ubiquitin moieties). Furthermore, activation by K63-linked ubiquitin increased with increasing chain length (Figure 3C and Figure 3D for quantification). Even though the UBZ domain promotes SPRTN binding to ubiquitin chains in pulldown assays (Ref^34,35^ and Figure S4A-C), it – and SPRTN’s MIU – was not required for activation (Figure 3E and Figure 3F for quantification). In contrast, Leu99 in SPRTN’s USD interface was strictly required for activation, while replacement of Leu38 had no apparent effect (Figure 3E and Figure 3F for quantification), which is consistent with its smaller predicted contribution to ubiquitin binding (Figure 1Q). Of note, the L99S substitution had no effect on ubiquitin chain binding in pull down assays (Figure S4A-C), suggesting that while the ubiquitin-USD interaction is critical for activation, it is too weak to withstand the washing steps during pulldown experiments.

**Figure 3.**
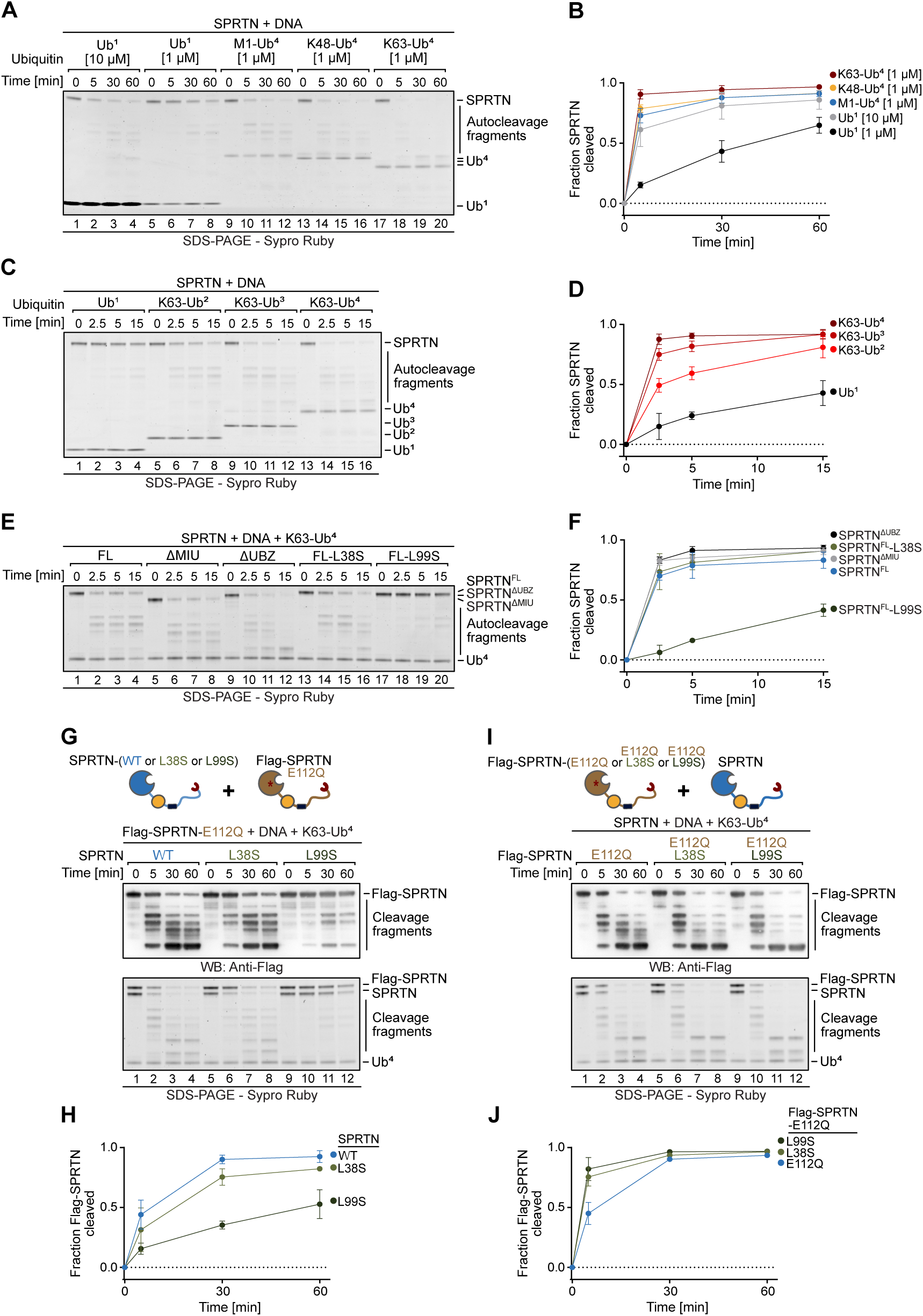
Poly-ubiquitin is particularly potent at SPRTN activation. (A) SPRTN^FL^ (250 nM) was incubated in the presence of Virion DNA (11.14 nM) with either mono-ubiquitin (Ub^1^) (1 or 10 µM), M1-tetra-ubiquitin (Ub^4^), K48-Ub^4^ or K63-Ub^4^ (1 µM). Ubiquitin concentrations refer to the concentration of individual ubiquitin moieties. (B) Quantification of SPRTN autocleavage assay shown in (A), depicting the mean ± SD of three independent experiments. (C) Reactions as in (A) but using Ub^1^, K63-Ub^2^, K63-Ub^3^ or K63-Ub^4^ (1 µM; referring to the concentration of individual ubiquitin moieties). (D) Quantification of SPRTN autocleavage assay shown in (C), depicting the mean ± SD of three independent experiments. (E) Reactions as in (A) but using the indicated SPRTN mutants and K63-Ub^4^. (F) Quantification of SPRTN autocleavage assay shown in (E), depicting the mean ± SD of four independent experiments. (G) Indicated SPRTN variants (250 nM) were incubated in the presence of Virion DNA (11.14 nM), Flag-SPRTN-E112Q (250 nM) and K63-Ub^4^ (1 µM, referring to the concentration of individual ubiquitin moieties). (H) Quantification of SPRTN autocleavage assay shown in (G), depicting cleavage of Flag-SPRTN-E112Q as the mean ± SD of three independent experiments. (I) SPRTN (250 nM) was incubated in the presence of single-stranded ΦX174 Virion DNA (11.14 nM), indicated Flag-SPRTN-E112Q variants (250 nM) and K63-Ub^4^ (1 µM referring to the concentration of individual ubiquitin moieties). Ubiquitin concentrations refer to the concentration of individual ubiquitin moieties. (J) Quantification of SPRTN autocleavage assay shown in (I). depicting cleavage of Flag-SPRTN-E112Q as the mean ± SD of three independent experiments. See also Figure S4.

Given that SPRTN autocleavage occurs *in trans*^26^, we considered the possibility that ubiquitin chains are particularly potent at promoting autocleavage because they can bridge two SPRTN molecules. To test this hypothesis, we assessed *in trans* cleavage of inactive SPRTN (Flag-tagged SPRTN^FL^-E112Q) by different active SPRTN variants (WT, L38S, or L99S). The WT enzyme cleaved the substrate efficiently in the presence of DNA and tetra-ubiquitin, while the L38S and the L99S variants showed a slight or strong reduction in activity, respectively (Figure 3G and Figure 3H for quantification). In the reverse scenario, we replaced Leu38 and Leu99 in inactive Flag-tagged SPRTN-E112Q and tested whether this affects cleavage by active WT SPRTN. In both cases, cleavage was at least as efficient as cleavage of inactive SPRTN with an intact USD interface (Figure 3I and Figure 3J for quantification). Together, these results show that the USD is dispensable in the SPRTN molecule that is cleaved but is required in the cleaving SPRTN molecule. Hence, ubiquitin chains stimulate autocleavage by activating one SPRTN molecule rather than bridging substrate and enzyme.

### Ubiquitylation of DNA-protein crosslinks promotes their cleavage by SPRTN

Having established that ubiquitin chains stimulate SPRTN autocleavage, we asked next whether the ubiquitylation of DPCs^10,15–21^ can trigger their cleavage by activating SPRTN. In a first instance, we tested whether the addition of free ubiquitin chains promotes the cleavage of DPCs formed *in vitro* between the catalytic SRAP domain of HMCES (HMCES^SRAP^) and an AP site-containing fluorescently-labeled ssDNA-dsDNA junction (Ref^44,45^ and Figure S5A-B). HMCES^SRAP^-DPCs were incubated with increasing concentrations of SPRTN and the helicase FANCJ. FANCJ is essential for DPC cleavage in these assays, because it unfolds the protein adduct by loading on the ssDNA 5’ of the DPC, then translocating into the crosslinked protein, resulting in its unfolding^44^. Upon unfolding, SPRTN can access the underlying DNA junction and cleave the protein adduct. As expected, DPC cleavage increased with higher SPRTN concentrations (Figure 4A, lane 3-5 and lanes 23-25) and required the presence of FANCJ (Figure 4A, lane 6 and lane 26 and Figure S5C, lane 3 vs 6). The need for FANCJ could be bypassed by thermal unfolding of the protein adduct, as observed previously (Ref^44^ and Figure S5C, lane 10). Upon addition of K48- or K63-linked tetra-ubiquitin chains, DPC cleavage was enhanced (Figure 4A, compare lanes 3-5 with lanes 7-9 (K48) and lanes 23-25 with 27-29 (K63)). Remarkably, in addition to the standard HMCES^SRAP^ cleavage fragment produced by SPRTN (cleaved DPC), additional smaller cleavage products (cleaved DPC*) appeared specifically in the presence of both types of ubiquitin chains. The appearance of these cleavage products was reduced (L38S) or absent (L99S) in SPRTN mutant variants with a non-functional USD interface (Figure S5D). These results demonstrate that ubiquitin chains stimulate DPC cleavage by interacting with SPRTN’s USD interface.

**Figure 4.**
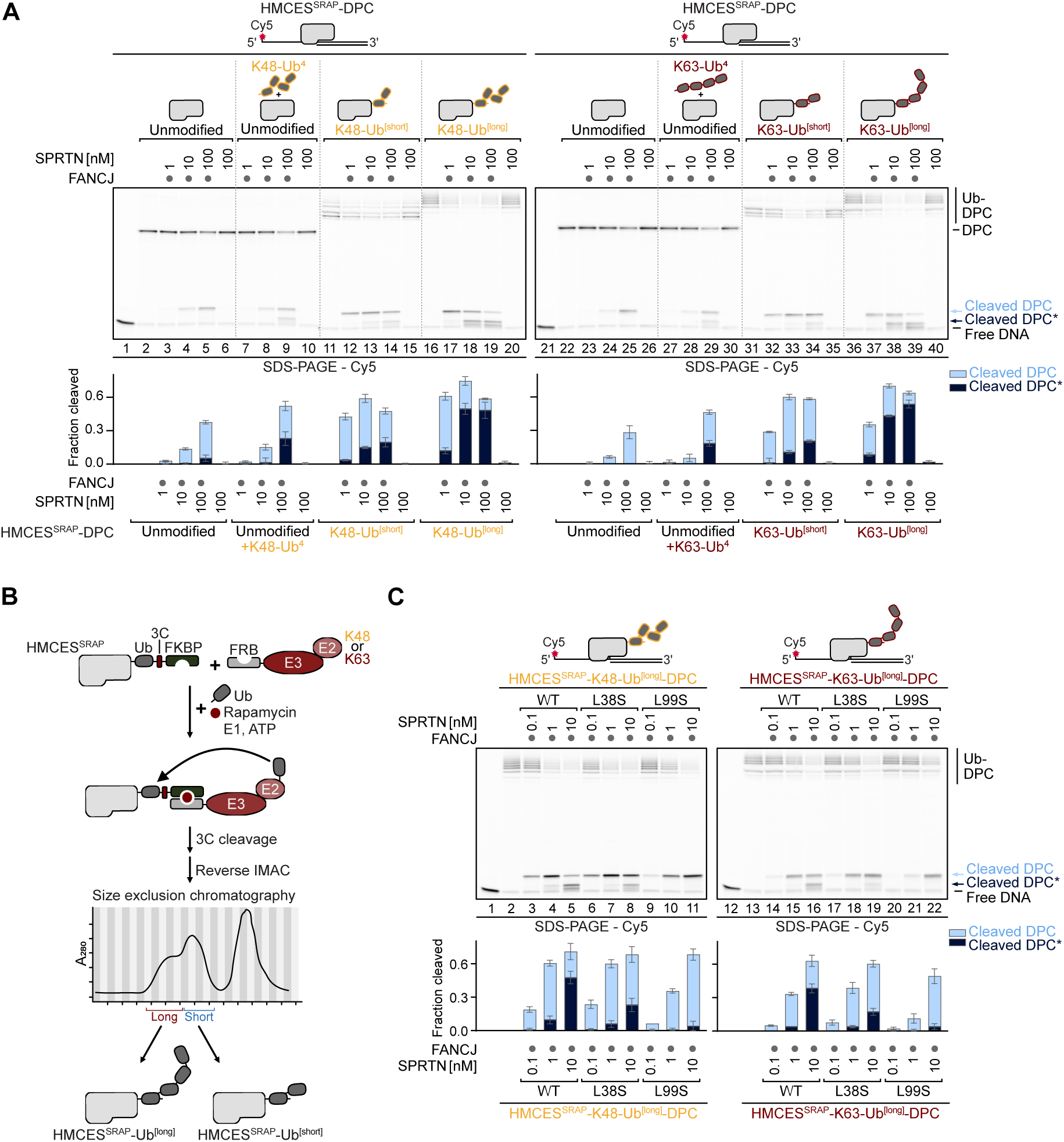
Ubiquitylation of DPCs promotes their cleavage by SPRTN. (A) Indicated HMCES^SRAP^-DPCs (10 nM) were incubated alone or in the presence of FANCJ (100 nM), SPRTN^FL^ (1-100 nM), K48-tetra-ubiquitin (Ub^4^) or K63-Ub^4^ (400 nM referring to the concentration of individual ubiquitin moieties) for 1 h at 30°C. Quantification: bar graphs represent the mean ± SD of three independent experiments. (B) Schematic of HMCES^SRAP^ ubiquitylation to generate DPCs shown in (A), (C) and Figure S6D. HMCES^SRAP^-Ub(G76V)-3C-FKBP was incubated with FRB-E3+E2 (K48 or K63) in the presence of ubiquitin, rapamycin, ubiquitin-E1 and ATP for 2 h (K63) or 6.5 h (K48) at 30°C. After cleavage of the FKBP-tag via 3C-protease, ubiquitylated HMCES^SRAP^ was purified by reverse immobilized metal affinity chromatography (IMAC) and size-exclusion chromatography (SEC). (C) Indicated HMCES^SRAP^-Ub^[long]^-DPCs (10 nM) were incubated alone or in the presence of FANCJ (100 nM) and SPRTN^FL^ (0.1-10 nM) for 1 h at 30°C. Quantification: bar graphs represent the mean ± SD of three independent experiments. See also Figure S5.

Motivated by these results, we set out to reconstitute DPC ubiquitylation *in vitro* to assess its effect on SPRTN activity. To this end, we employed synthetic engineered ubiquitin E3s (streamlined versions of the previously published Ubiquiton system^46^) to modify HMCES^SRAP^ with K48- or K63-linked ubiquitin chains prior to DPC formation (Figure 4B). First, we fused a *C*-terminal tag containing a mono-ubiquitin moiety, as well as an FK506-binding protein (FKBP) domain to HMCES^SRAP^. We then incubated this substrate with ubiquitin, an engineered ubiquitin E3 carrying an FKBP-rapamycin-binding (FRB) domain, ubiquitin activating enzyme (E1), ubiquitin conjugating enzyme (E2), ATP and rapamycin. Rapamycin induces proximity between the substrate and the E3, promoting modification of the single ubiquitin moiety fused to HMCES^SRAP^ with either K48- or K63-linked polyubiquitin chains (depending on the identity of the E2/E3 enzymes used in the assay). Following cleavage of the 3C site between ubiquitin and the FKBP domain, HMCES^SRAP^ modified with short (HMCES^SRAP^-Ub^[short]^) or long (HMCES^SRAP^-Ub^[long]^) ubiquitin chains was purified over several steps (Figure 4B and S5A).

DPC ubiquitylation strongly enhanced DPC cleavage by SPRTN, independently of linkage type, with long chains activating stronger than shorter ones (Figure 4A, lanes 11-20 (K48) and 31-40 (K63)). Of note, the extent of cleavage of ubiquitylated DPCs by 1 nM of SPRTN was comparable to the cleavage of unmodified DPCs by 100 nM of SPRTN (Figure 4A, compare lanes 5 and 17 (K48) and lanes 25 and 37 (K63)), indicating that DPC ubiquitylation activates SPRTN at least a hundred-fold. As observed above in the presence of free chains, DPC ubiquitylation led to the appearance of smaller DPC cleavage products (cleaved DPC*), especially at high SPRTN concentrations (Figure 4A, lanes 18-19 (K48) and lanes 38-39 (K63)). The USD interface was required for these effects. SPRTN-L38S and SPRTN-L99S mutant variants displayed reduced (L38S) or no (L99S) formation of these products (Figure 4C). These results show that DPC ubiquitylation allosterically activates SPRTN independently of linkage type by binding to its USD interface, enabling the protease to cleave the crosslinked protein more efficiently and completely.

### SUMO-targeted DPC ubiquitylation activates SPRTN in cells and *in vitro*

Next, we asked whether the activation of SPRTN by DPC ubiquitylation at the USD interface is required for SPRTN function in cells. SPRTN activation can be monitored in cells upon induction of DNA methyltransferase 1 (DNMT1)-DPCs using 5-azadC, a chemotherapeutic cytidine analog. 5-azadC is incorporated into DNA during replication, covalently entrapping DNMT1 in post-replicative chromatin^47^. Following DPC formation, the DNMT1 adduct is swiftly SUMOylated, which requires SUMO E3s of the PIAS family^22^, followed by ubiquitylation by SUMO-targeted ubiquitin E3s RNF4^10,15,16^ and TOPORS^17,18^. In a first instance, we thus depleted RNF4 using siRNA in HAP1 WT and *TOPORS* knock-out cells and assessed the effect on 5-azadC-induced SPRTN autocleavage. Simultaneous depletion of RNF4 and TOPORS, and thereby DPC ubiquitylation, resulted in a complete loss of SPRTN autocleavage fragments (Figure S6A), indicating that DPC ubiquitylation is critical for efficient SPRTN activation in cells.

To directly test whether DNMT1-DPC cleavage requires the ubiquitin-dependent activation of SPRTN, we complemented HeLa T-REx Flp-In cells that are genetically engineered to express patient-mimicking *SPRTN^ΔC^*alleles with doxycycline-inducible YFP-tagged SPRTN variants (WT, E112Q, L38S and L99S). While *SPRTN^ΔC^* cells are viable, they fail to efficiently cleave 5-azadC-induced DNMT1-DPCs^10^. To monitor DPC repair, we synchronized cells via double-thymidine block, induced expression of YFP-tagged SPRTN variants, released cells into early S-phase, and induced DNMT1-DPCs by 5-azadC treatment for 30 min, followed by an optional 2-h chase in drug-free media to allow repair (Figure 5A, top). DNMT1-DPC formation was assessed by purification of x-linked proteins (PxP) (Ref^10,48^, Figure 5A and Methods). DNMT1-DPCs were detected in all cell lines upon 5-azadC treatment (Figure 5A). Following the 2-h chase, a characteristic SPRTN cleavage fragment formed in *SPRTN^ΔC^* cells expressing SPRTN-WT but not in cells expressing catalytically inactive SPRTN-E112Q (Figure 5A, red dots), as observed previously^10^; DPCs are still resolved in these cells because crosslinked DNMT1 is additionally targeted by proteasomal degradation^10,15^. The SPRTN-dependent cleavage product was reduced in cells expressing SPRTN-L38S and almost undetectable in cells expressing SPRTN-L99S (Figure 5A, red dots), indicating that SUMO-targeted ubiquitylation activates SPRTN in cells to promote DPC cleavage.

**Figure 5.**
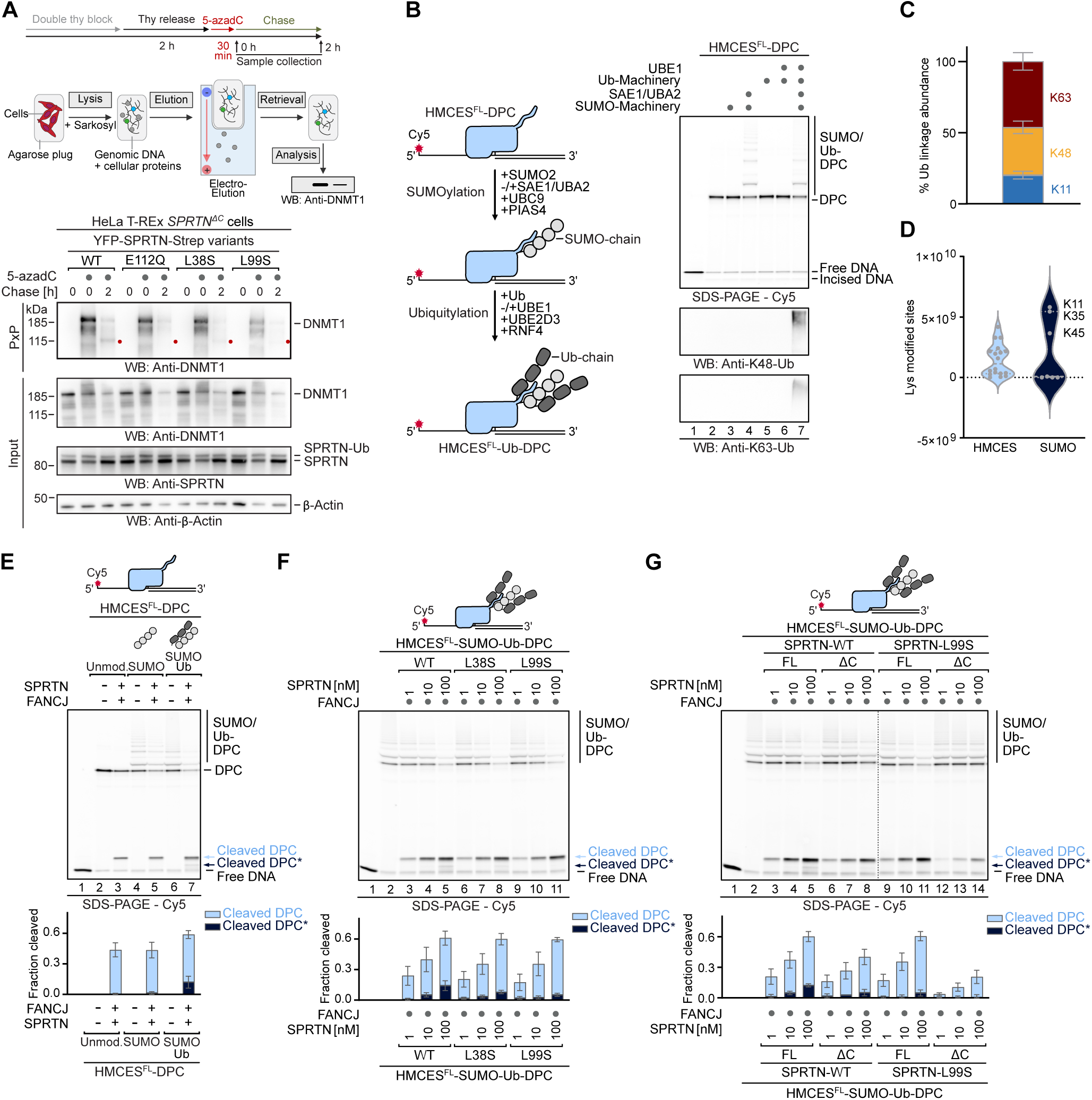
Replication-independent DPC repair by SPRTN is mediated by the USD. (A) HeLa-TREx *SPRTN^ΔC^* Flp-In complemented with indicated YFP-SPRTN^FL^-Strep-tag variants were treated as depicted (top) with 5-azadC (10 µM) and harvested at indicated time points. DNMT1-DPCs were isolated using PxP (middle, see Methods) and analyzed by immunoblotting (bottom). (B) Schematic of SUMO-targeted ubiquitylation of HMCES^FL^-DPCs used in (B-G), and Figure S6C (left). HMCES^FL^-DPCs were incubated alone or in the presence of SUMO2, UBC9 and PIAS4, with or without SAE1/UBA2 for 30 min. Next unmodified or SUMOylated HMCES^FL^-DPCs were incubated alone or in the presence of ubiquitin (Ub), RNF4, UBE2D3, with or without UBE1 for 30 min, prior to separation by denaturing SDS-PAGE and immunoblotting (right). (C) Mass spectrometry analysis of ubiquitin linkages formed SUMO-targeted ubiquitylation of HMCES^FL^-DPCs. Bar chart shows the mean ± SD of four biological replicates. (D) Mass spectrometry analysis of lysine residues within HMCES or SUMO modified upon SUMO-targeted ubiquitylation. Violin blots show the mean ± SD of four biological replicates. (E-G) Indicated HMCES^FL^-DPCs (10 nM) were incubated alone or in the presence of FANCJ (100 nM) and the indicated concentrations of SPRTN (WT or mutant variants, as indicated) for 1 h at 30°C. Quantifications: bar graphs represent the mean ± SD of three independent experiments. Values for SPRTN^FL^ were also used for quantification of Figure S6C. See also Figure S6.

Encouraged by these results, we wanted to reconstitute the SUMO-dependent ubiquitylation of DPCs *in vitro*. Therefore, we generated DPCs between HMCES and a Cy5-labeled ssDNA-dsDNA junction (Figure S6B)^44,45^; we used full-length HMCES (HMCES^FL^) because it contains a canonical SUMOylation site in its *C*-terminal tail that is absent in HMCES^SRAP^ constructs. HMCES-DPCs were incubated with the SUMOylation machinery, consisting of the SUMO-E1 SAE1/UBA2, the SUMO-E2 UBC9, the SUMO-E3 PIAS4, SUMO2 and ATP (Figure S6B). The successful SUMOylation of the crosslinked protein was indicated by slower migrating species of the HMCES-DPCs (Figure 5B, lane 4). No SUMOylation was observed in reactions lacking SUMO-E1 (Figure 5B, lanes 3). For the subsequent ubiquitylation, SUMOylated DPCs were incubated with ubiquitin, ubiquitin-E1, ubiquitin-E2 UBE2D3 and the SUMO-targeted ubiquitin-E3 RNF4 (Figure S6B). Ubiquitylation of SUMOylated DPCs was evident as further upshifts in gel migration and was in addition confirmed by western blot (Figure 5B, lane 7). To determine the identity of the ubiquitylated lysine residues and the involved ubiquitin linkages, we used mass spectrometry (MS). We identified K48-, K63- and K11-linked ubiquitin chains on SUMOylated DPCs (Figure 5C), as has been observed in cells^18^. Ubiquitin chains were formed on various lysine residues of HMCES and on three distinct lysine residues of SUMO2 (Figure 5D). Ubiquitylation was lost in the absence of ubiquitin-E1 or in the absence of SUMOylation (Figure 5B, lanes 5 and 6), demonstrating *bona fide* SUMO-targeted DPC ubiquitylation.

Upon incubation with SPRTN, we observed enhanced cleavage of the ubiquitylated protein adduct, compared to unmodified DPCs and SUMOylated DPCs (Figure 5E, compare lane 3 and 5 with lane 7). While we again observed the appearance of additional DPC cleavage products (Figure 5E, lane 7, Cleaved DPC*), they were less abundant than in experiments using the synthetic ubiquitylation system (Figure 4), presumably because not all DPCs are modified by SUMO-targeted ubiquitylation in our experiments. The additional cleavage products were reduced (L38S) or absent (L99S) when DPCs were degraded with SPRTN variants with a mutated USD interface (Figure 5F, compare lanes 3-5 with lanes 6-8 and 9-11).

The repair of DNMT1-DPCs in cells is not only compromised upon replacement of critical USD residues (Figure 5A), but also upon loss of SPRTN’s *C*-terminal tail in RJALS SPRTN^ΔC^ patient variants^10^. Therefore, we tested the ability of SPRTN^ΔC^ to process DPCs modified by SUMO-targeted ubiquitylation *in vitro*. While SPRTN^ΔC^ displayed slightly reduced DPC cleavage compared to the WT enzyme (Figure 5G, compare lanes 3-5 with lanes 6-8), the extent of cleavage was strongly reduced upon additional replacement of Leu99 by serine (SPRTN^ΔC^-L99S) (Figure 5G, compare lanes 6-8 and lanes 12-14). The synthetic cleavage defect of SPRTN^ΔC^-L99S was only partially explained by loss of the UBZ domain, given that SPRTN^ΔUBZ^-L99S variant displayed a less pronounced phenotype (Figure S6C, lanes 12-14). Notably, the defect of SPRTN^ΔC^ was specific to DPCs modified by SUMO-targeted ubiquitylation. DPCs modified on their *C*-terminus using the synthetic ubiquitylation system were cleaved comparably well by SPRTN^ΔC^ and the WT enzyme (Figure S6D, compare lanes 3-5 with lanes 6-8 (K48) and compare lanes 17-19 with lanes 20-22 (K63)), while a USD mutant variant (L99S) displayed clear defects (Figure S6D, lanes 9-14 (K48) and 23-28 (K63)). Taken together, our results suggest that the SUMO-targeted ubiquitylation of DPCs ensures efficient repair by allosteric activation of SPRTN at the USD interface. *In vitro*, the ubiquitin-dependent activation of SPRTN appears to be specifically important to support the residual cleavage of RJALS SPRTN^ΔC^ patient variants towards DPCs modified by SUMO-targeted ubiquitylation (see Discussion).

### Ubiquitin-dependent activation of SPRTN maintains genome stability in Ruijs-Aalfs syndrome

Next, we wanted to determine whether the ubiquitin-dependent activation of SPRTN at the USD interface is also essential for the function of SPRTN^ΔC^ patient variants in cells. To this end, we complemented conditional *Sprtn^F/-^ CreER^T2^* knock-out mouse embryonic fibroblasts (MEFs) with either an empty vector (EV) or different human SPRTN variants (FL and ΔC) tagged with a *C*-terminal Strep-tag (Figure S7A-B). Of note, SPRTN^ΔC^ variants express at much higher levels than the WT enzyme, which is consistent with observations in RJALS patients (Figure S7A-B)^38^. Knock-out was induced upon addition of 4-hydroxytamoxifen (4-OHT), with the solvent MeOH serving as control (Figure S7C-D), resulting in diverse phenotypes including growth arrest (Figure 6A-B), formation of micronuclei and chromatin bridges (Figure 6C-E), as wells as arrest in the G2/M phase of the cell cycle (Figure S7E-F), as described previously^29^. All phenotypes were rescued by expression of human WT SPRTN but not by catalytically inactive SPRTN-E112Q (Figure 6A and 6D). Also, expression of SPRTN^ΔC^ complemented all phenotypes induced by *Sprtn* knock-out (Figure 6B and 6E). While the replacement of USD residues Leu38 or Leu99 had no effect on the ability of SPRTN^FL^ to complement cell fitness and cell cycle defects upon loss of mouse *Sprtn* (Figure 6A and Figure S7E), loss of Leu99 resulted in intermediate growth defects and G2/M arrest in SPRTN^ΔC^ (Figure 6B and Figure S7F). These defects were accompanied by severe signs of genome instability, observed as micronuclei and chromatin bridges in cells expressing SPRTN^ΔC^-L99S (Figure 6C+E) and to a lesser degree for SPRTN^ΔC^-L38S (Figure 6E).

**Figure 6.**
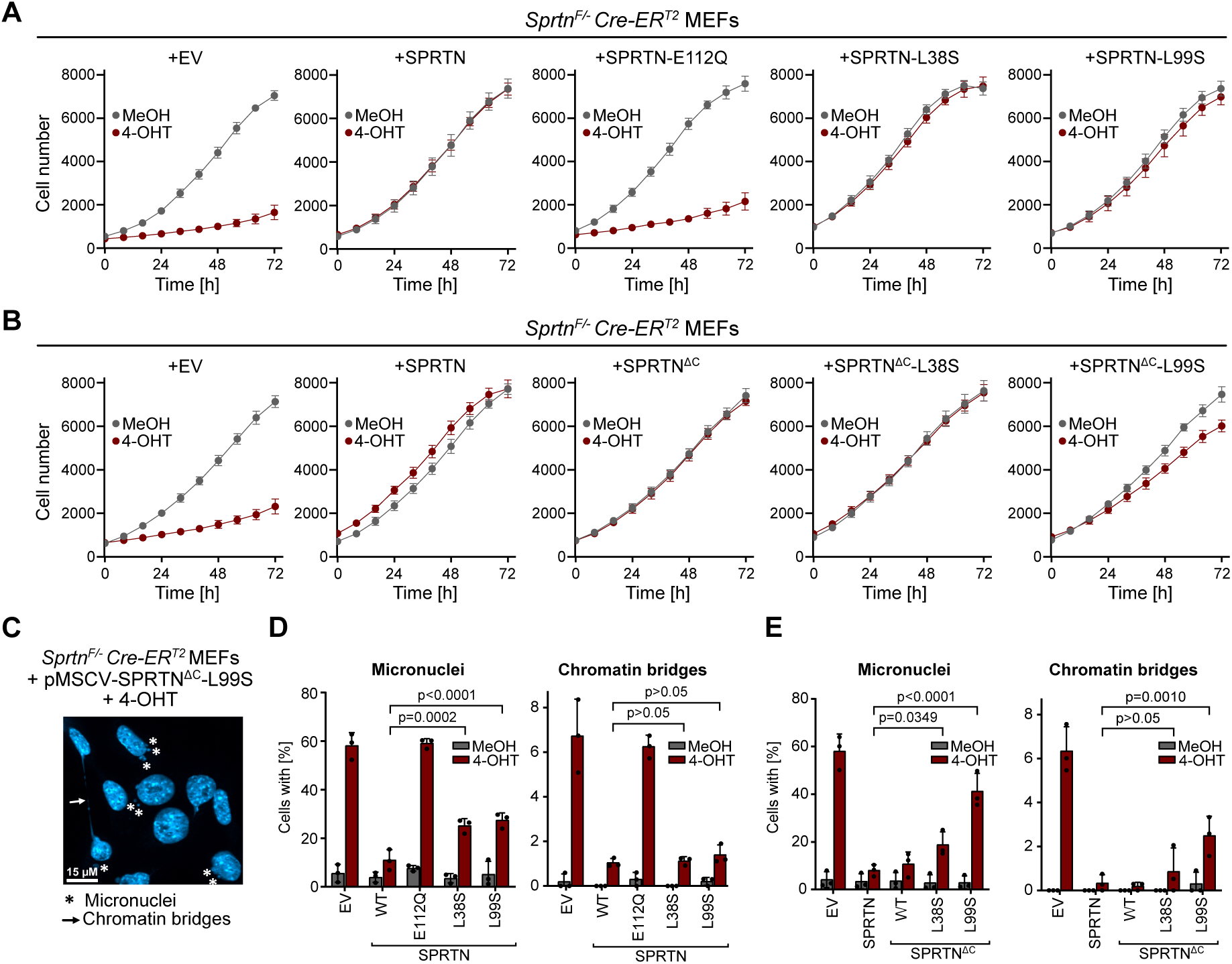
Ubiquitin-dependent activation of SPRTN maintains genome stability in Ruijs-Aalfs syndrome. (A-B) Proliferation of *Sprtn^F/-^* Cre-ER^T2^ mouse embryonic fibroblasts (MEFs) complemented with indicated SPRTN variants or empty vector (EV, pMSCV) treated with methanol (MeOH) or (Z)-4-hydroxytamoxifen (4-OHT) (2 µM) for 48 h. After seeding, cell numbers were counted at indicated time points. Values are the mean ± SD of eight technical replicates. Shown is a representative of three independent experiments. (C) Image showing micronuclei (asteriks) and chromatin bridges (arrow) in *Sprtn^F/-^*Cre-ER^T2^ MEFs + pMSCV-SPRTN^ΔC^-L99S treated with 4-OHT (2 µM) for 48 h. DNA was visualized by DAPI staining. Scale bar corresponds to 15 µM. (D-E) Quantification of micronuclei and chromatin bridges formation in *Sprtn^F/-^* Cre-ER^T2^ MEFs complemented with indicated SPRTN variants or EV (pMSCV) treated with MeOH or 4-OHT (2 µM) for 48 h. DNA was visualized by DAPI staining. Bar graphs show the mean ± SD of three independent experiments. The p values were calculated using a two-way ANOVA with Dunnett’s multiple comparison test. See also Figure S7.

Collectively, these experiments demonstrate that SPRTN’s USD interface, and thus the allosteric activation of SPRTN by ubiquitin, is critical to maintain fitness and genome stability in cells expressing truncated RJALS patient variants.

## DISCUSSION

Over the last decade, DPC repair has emerged as a conserved cellular process that is essential for maintaining genome stability^5^. Since the identification of dedicated DPC proteases in yeast and humans^11,24,26,27,49–52^, it has remained enigmatic how specificity for crosslinked protein adducts is achieved. The DPC protease SPRTN features a bipartite DNA binding module, consisting of ZBD and BR, which provides a first layer of specificity by restricting activity to the cleavage of DPCs near ssDNA-dsDNA junctions and other structures with single- and double-stranded features^19,31,32^. However, such junctions are widespread across the genome. Hence, SPRTN’s DNA structure-specific activity alone is insufficient to explain how the protease achieves specificity.

Our study reveals that SPRTN activation is linked to the ubiquitylation of the crosslinked protein. By reconstituting DPC ubiquitylation *in vitro*, we observed that this modification stimulates SPRTN activity by up to two orders of magnitude, regardless of ubiquitin chain linkage type (Figure 4A). Ubiquitin activates SPRTN by binding to the USD interface at the back of the protease domain (Figure 1N and 2A). The ubiquitin-dependent activation allows the enzyme to process DPCs not only more efficiently but also to a greater extent, which may be crucial for enabling efficient translesion synthesis bypass of the remaining peptide adduct during replication-coupled DPC repair^11^. Our MD simulations further suggest that the USD-ubiquitin interaction stabilizes an open conformation of the enzyme, exposing its active site. This model is supported by our NMR data, which additionally indicate that the ubiquitin-induced change of SPRTN’s conformation is especially pronounced in the presence of an activating DNA structure (Figure 2A). Indeed, while ubiquitin enhances SPRTN’s DNA-dependent activity, it is not sufficient to trigger SPRTN activation by itself (Figure 1B).

These insights help explain why cells extensively ubiquitylate DPCs^10,11,13–21^ and why ubiquitylated DPCs accumulate in cells following SPRTN depletion^39^. The observation that SPRTN-dependent cleavage can occur without DPC ubiquitylation in frog egg extracts^19,20^ is not necessarily inconsistent with our findings. It is plausible that the DPC cleavage observed in egg extract in the absence of ubiquitylation originates from SPRTN’s basal, ubiquitin-independent activity, which is also evident in our assays.

Consistently, while amino acid substitutions within the USD interface substantially reduced DPC cleavage activity *in vitro* and in cells, they did not completely abolish SPRTN function. SPRTN with a replacement of the USD residue Leu99, which showed stronger phenotypes across all our assay compared to replacing Leu38, was able to generally complement the loss of *Sprtn* in MEFs. The same is true for the RJALS SPRTN^ΔC^ patient variant, which also displayed strongly reduced DNMT1-DPC cleavage^10^, while successfully complementing *Sprtn* loss. Thus, only a minimal amount of SPRTN activity appears to be necessary to fulfil its essential role in suppressing genome instability. The critical role of the USD became evident when Leu99 was replaced in SPRTN^ΔC^, resulting in cell fitness defects and formation of micronuclei and chromatin bridges in mitosis (Figure 6).

The synthetic effect observed between the combined loss of SPRTN’s *C*-terminal tail and a functional USD interface, is only partially explained by the loss of the UBZ domain. While the UBZ is required for efficient DPC cleavage in cells^10^ and egg extracts^19^, it showed weaker defects in processing ubiquitylated DPCs in combination with the L99S substitution than the corresponding SPRTN^ΔC^-L99S variant (Figure 5G and Figure S6C). In addition to lacking the UBZ domain, SPRTN^ΔC^ also exhibits reduced DNA binding compared to the WT protein^26,27^, which may contribute to its reliance on the USD for full functionality. Based on these considerations, we propose a partially speculative ‘triple-lock’ model in which SPRTN activity is controlled by at least three mechanisms (Figure 7). First, the UBZ supports SPRTN function by recruiting it to ubiquitylated DPCs, as previously suggested^10,19^. This recruitment function is likely more important in the crowded environment of a cell than in our *in vitro* experiments, explaining why the loss of the UBZ had no or only weak phenotypes in our assays. Second, the binding of an activating DNA structure induces an open conformation of SPRTN. Third, this open, active conformation is further stabilized by binding of ubiquitylated DPCs to SPRTN’s USD interface, facilitating rapid and complete proteolysis of the crosslinked protein adduct.

**Figure 7.**
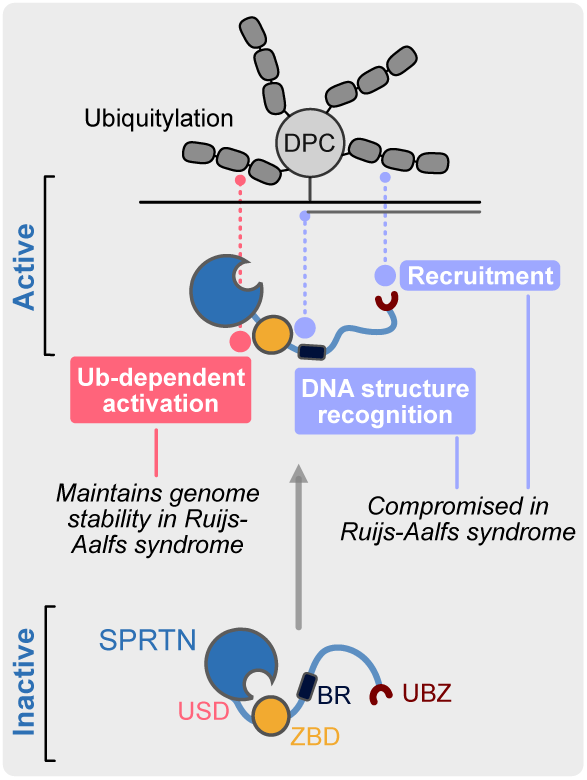
‘Triple-lock’ model for SPRTN activation. The ubiquitin-binding zinc finger (UBZ) recruits SPRTN to ubiquitylated DPCs. Binding of both DNA-binding domains, zinc-binding domain (ZBD) and the basic region (BR) to activating DNA structures induces an open conformation. This open conformation is stabilized by ubiquitin binding to the ubiquitin binding interface at the SprT domain (USD). Recruitment and DNA structure recognition are compromised in Ruijs-Aalfs syndrome patients, which therefore fully rely on the Ub-dependent activation via the USD to maintain genome stability.

This model offers a potential explanation for why SPRTN^ΔC^ displayed defects in processing DPCs modified by SUMO-targeted ubiquitylation (Figure 5G) but not of DPCs modified using the artificial ubiquitylation system (Figure S6D). In the synthetic system, the DPC is modified exclusively at the *C*-terminal ubiquitin tag. In contrast, our MS analysis revealed that SUMO-targeted ubiquitylation affects multiple lysine residues within the DPC and the SUMO chain (Figure 5C-D). Some of these ubiquitylation events may hinder SPRTN function by interfering with efficient DNA binding. Thus, SPRTN’s full DNA binding capacity is likely required in this context.

The ability of SPRTN to be activated by both K48- and K63-linked ubiquitin chains suggests a first hypothesis as to why SPRTN is essential, despite acting redundantly with the proteasome in most experimental systems investigated so far^10,19,20^. The proteasome mainly targets substrates modified with K48-linked ubiquitin^53^. Therefore, our data implies that key endogenous substrates of SPRTN may be modified by K63-linked ubiquitin and are consequently not amenable to proteasomal degradation.

In conclusion, we have demonstrated how the key DNA repair enzyme SPRTN is precisely deployed. The recruitment of DNA repair factors via canonical ubiquitin binding domains is a common theme in the DNA damage response. However, ubiquitin does not only recruit SPRTN but also allosterically activates the enzyme, which is crucial for maintaining genome stability in RJALS patients.

## MATERIALS AND METHODS

### Cell lines

#### Insect cell lines

High Five cells (Invitrogen) were cultured at 27°C for protein overexpression. Cells were cultured in ESF 921 insect cell culture medium (Thermo Scientific).

#### Mammalian cell lines

HeLa TREx Flp-In *SPRTN^ΔC^* cells (Ref^10^) stably expressing YFP-SPRTN-Strep-tag variants were generated using the Flp-In system (pOG44, Thermo Scientific) according to manufacturer’s instructions and selected in Hygromycin B (150 µg/mL) (Thermo Scientific). Protein expression was induced by overnight incubation with doxycycline (DOX) (Sigma) (final concentration 1 µg/mL). HeLa TREx Flp-In *SPRTN^ΔC^* cells were grown in Dulbecco’s Modified Eagle Medium (DMEM) supplemented with 10% (v/v) fetal bovine serum (FBS).

HAP1 wildtype (WT) cells and HAP1 *TOPORS* knock-out (KO) cells were grown in Iscove’s Modified Dulbecco’s Medium (IMDM) supplemented with 10% (v/v) FBS and 1% (v/v) Penicillin-Streptomycin-Glutamine (PSG).

*Sprtn^F/-^* mouse embryonic fibroblasts (MEFs) (H7) immortalized with SV40 large T and transduced with a retroviral vector expressing Cre-ER^T2^ (Ref^29^) were cultured in DMEM supplemented with 10% (v/v) FBS. *Sprtn* KO was induced by treating 4×10^5^ cells with methanol (MeOH) (vehicle control) or 2 μM (Z)-4-hydroxytamoxifen (4-OHT) (Sigma) for 48 h. Conversion of the floxed *Sprtn* allele (F) to the KO allele (-) was verified by PCR using WT- and KO-specific forward primers and a common reverse primer. PCR conditions were 35 cycles of 94°C for 30 s, 60°C for 30 s, and 72°C for 1 min. PCR products are 527 bp and 278 bp for the floxed and the KO alleles respectively. For exogenous expression of human SPRTN in MEFs, cells were infected with retroviral vectors produced in HEK293T/17 (ATCC, CRL-11268) by co-transfecting pMSCV.hyg-SPRTN-Strep with gag-pol and VSV-G packaging plasmids. Infected cells were selected with Hygromycin B (200 µg/mL) (Thermo Scientific) for 8 days.

### Protein expression and purification

#### SPRTN

Recombinant (Flag-tagged) SPRTN (Full-length, ΔMIU, and ΔUBZ – WT or in combination with L38S, L99S, and E112Q amino acid replacements) protein was expressed in BL21 (DE3) *E. coli* cells and purified as previously described with slight modifications^32^.

BL21 (DE3) *E. coli* cells were grown at 37°C in Terrific broth (TB) medium (prepared with tap water) until they reached OD 0.7. Protein expression was induced by addition of 0.5 mM Isopropyl-β-D-thiogalactoside (IPTG) overnight at 18°C. The next day, cells were harvested, snap-frozen in liquid nitrogen and stored at −80°C. All subsequent steps were carried out at 4°C. For protein purification, cell pellets were resuspended in buffer A (50 mM HEPES/KOH pH 7.2, 500 mM KCl, 1 mM MgCl_2_, 10% Glycerol, 0.1% IGEPAL, 0.04 mg/mL Pefabloc SC, cOmplete EDTA-free protease inhibitor cocktail tablets, 1 mM Tris(2-carboxyethyl)phosphine hydrochloride (TCEP)) and lysed by sonication. Cell lysate was incubated with smDNAse (45 U/mL lysate) for 30 min on a roller prior to removal of cell debris by centrifugation at 18,000 g for 30 min. Cleared supernatant was filtered using syringe filters (PVDF, 0.22 µm) and applied to Strep-Tactin®XT 4Flow® high-capacity cartridges, washed with 3 column volumes (CV) of buffer A and 4 CV of buffer B (50 mM HEPES/KOH pH 7.2, 500 mM KCl, 10% Glycerol, 1 mM TCEP). Proteins were eluted in 6 CV buffer C (50 mM HEPES/KOH pH 7.2, 500 mM KCl, 10% Glycerol, 1 mM TCEP and 50 mM Biotin). Eluted proteins were further applied to HiTrap Heparin HP affinity columns and washed with 3 CV buffer B before eluting in buffer D (50 mM HEPES/KOH pH 7.2, 1 M KCl, 10% Glycerol, 1 mM TCEP). Eluted fractions containing recombinant SPRTN protein were desalted against buffer B using PD-10 desalting columns. The affinity tag was cleaved off at 4°C overnight by addition of His-tagged TEV protease with 1:10 mass ratio. Cleaved recombinant SPRTN protein was further purified by size exclusion chromatography (SEC) using a HiLoad 16/600 Superdex 200 pg column equilibrated in buffer B (50 mM HEPES/KOH pH 7.2, 500 mM KCl, 10% Glycerol, 1 mM TCEP). Eluted proteins were concentrated with 10 kDa cutoff Amicon Ultra centrifugal filters before aliquoting, snap-freezing in liquid nitrogen and storage at −80°C.

Following SPRTN purification, metalation of the protein was examined by Inductively Coupled Plasma Optical Emission Spectrometry (ICP-OES), which confirmed correct metalation with three Zn^2+^ ions per full-length SPRTN molecule.

For truncated SPRTN variants smaller than 30 kDa including SPRTN^ΔC^ (WT or L99S), SprT-BR (WT, E122Q, E112Q+L99S), ZBD-BR and ZBD, a Strep-tagged TEV protease was used. Prior to SEC, Strep-tagged TEV protease, residual uncleaved protein and the cleaved Tag were removed by a Strep-Tactin®XT 4Flow® high capacity cartridges^10^.

For NMR experiments, SprT-BR (E112Q, E112Q+L99S), ZBD, and ZBD-BR were expressed in ^15^N- or ^13^C-/^15^N-containing media. Here, cells were grown to OD 0.4, before the temperature of the incubator was lowered to 18°C and MnCl_2_ was added to a final concentration of 1.5 mM. Once OD 0.7 was reached expression was induced with 0.5 mM IPTG and performed overnight at 18°C. For SEC, buffer E (50 mM HEPES/KOH pH 7.2, 500 mM KCl, 1% Glycerol, 2 mM TCEP, pH 7.2) was used.

#### Mono-Ubiquitin

For purification of mono-ubiquitin (Ub^1^) a plasmid encoding Ub^1^ with a *N*-terminal His6-Tag was provided by Brenda Schulman (MPI for Biochemistry, Martinsried, Germany). Ub^1^-I44A was generated by introducing point mutations using the Q5-site-directed mutagenesis kit (NEB). Protein was expressed in Rosetta *E. Coli* cells, grown at 37°C in TB (prepared with tap water) to OD 0.7. Protein expression was induced with 0.5 mM IPTG overnight at 18°C. Cells were harvested the next day and directly resuspended in buffer A (50 mM Tris-HCl pH 7.5, 250 mM NaCl) (20 mL/ L culture), snap-frozen in liquid nitrogen and stored at −80°C. All subsequent steps were carried out at 4°C. For protein purification, cell lysates were thawed and Pefabloc SC (0.04 mg/mL), MgCl_2_ (1 mM) and smDNAse (45 U/mL lysate) were added. Cells were lysed by sonication and incubated for 30 min on a roller prior to removal of cell debris by centrifugation at 50,000 g for 30 min. Clarified lysate was filtered using syringe filters (PVDF, 0.22 µm) and mixed with Ni-NTA Agarose (QIAGEN) equilibrated in buffer A and incubated for 1 h on a roller to allow binding. The beads were transferred to a gravity flow column, washed with 15 CV of buffer A and protein was eluted in fractions of 1 CV each with buffer B (50 mM Tris-HCl pH 7.5, 250 mM NaCl, 300 mM imidazole). Fractions were checked via SDS-PAGE and Coomassie-based staining for presence of Ub^1^. Ub^1^-containing fractions were pooled and after addition of GST-tagged 3C-protease (0.5 mg/L culture), dialyzed against buffer A overnight. Cleaved protein was passed through Ni-NTA Agarose (QIAGEN) the next day for removal of uncleaved protein and His6-tag. The flow-through was collected, concentrated to 1 mL and loaded on a Superdex 200 Increase 10/300 GL column equilibrated in buffer C (50 mM HEPES/KOH pH 7.2, 500 mM KCl, 5% Glycerol, 2 mM TCEP). For Ub^1^ used for NMR experiments, this buffer was slightly modified, buffer C-NMR (50 mM HEPES/KOH pH 7.2, 500 mM KCl, 1% Glycerol, 2 mM TCEP). Eluted protein was concentrated with 10 kDa cut-off Amicon ultra centrifugal filters before aliquoting, snap-freezing in liquid nitrogen and storage at −80°C.

#### FANCJ

Recombinant FANCJ protein was expressed in High Five cells and purified as previously described^45^.

#### HMCES^SRAP^

Recombinant HMCES^SRAP^ (-WT, -R98E), protein was expressed in BL21 (DE3) *E. coli* cells and purified as previously described^45^, using TB prepared with tap water. For synthetic ubiquitylation of HMCES^SRAP^, a sequence encoding for Ub^1^(G76V) followed by an FKBP-domain, including linkers and a 3C protease cleavage site was codon optimized for bacterial expression and inserted at the *C*-terminal end of HMCES^SRAP^, in front of the His6-tag, in the pNIC_HMCES^SRAP^ plasmid. Purification followed protocols described for HMCES^SRAP^ and the final protein was further processed as described below.

#### HMCES^FL^

Recombinant HMCES^FL^ protein was expressed in BL21 (DE3) *E. coli* cells and purified as previously described^45^, analogously to recombinant SPRTN using TB prepared with tap water.

#### UBC9

For purification of recombinant UBC9, the open reading frame was codon optimized and cloned in a pBAD plasmid carrying a *N*-terminal His6-tag. Protein was expressed in BL21(DE3) *E. coli* cells and grown in TB (prepared with tap water) at 37°C to OD 0.7 before induction of protein expression with 0.1% L-arabinose at 18°C overnight. Cells were harvested the next day, snap-frozen in liquid nitrogen and stored at −80°C. All subsequent steps were performed at 4°C. For protein purification, cell pellets were thawed, resuspended in buffer A (20 mM HEPES/KOH pH 7.0, 2 mM Mg(OAc)_2_, 300 mM KOAc, 10% glycerol, 30 mM imidazole, 0.1% IGEPAL, 1 mM TCEP, cOmplete protease inhibitor, 0.04 mg/mL Pefabloc SC, 1 mg/mL lysozyme, 45 U/mL smDNAse) and incubated on a roller for 30 min. The lysate was sonicated for 15 min prior to cell debris removal by centrifugation at 18,000 g for 40 min. Clarified lysate was filtered using syringe filters (PVDF, 0.22 µm) and incubated with Ni-NTA agarose (QIAGEN) on a roller for 1 h at 4°C. The beads were transferred to a gravity flow column and washed with 15 CV buffer B (20 mM HEPES/KOH pH 7.0, 2 mM Mg(OAc)_2_, 300 mM KOAc, 10% glycerol, 30 mM imidazole) before elution in 2 CV buffer C (20 mM HEPES/KOH pH 7.0, 2 mM Mg(OAc)_2_, 300 mM KOAc, 10% glycerol, 300 mM imidazole). The His6-tag was cleaved by the addition of His-tagged TEV protease (1 mg/L culture) and dialyzed overnight against buffer D (20 mM HEPES/KOH pH 7.0, 2 mM Mg(OAc)_2_, 300 mM KOAc). The next day, cleaved protein was passed through Ni-NTA agarose to remove His-tagged TEV protease, residual uncleaved protein and His6-Tag. Flow-through was concentrated to 1 mL and loaded on a Superdex 200 Increase 10/300 GL column equilibrated in buffer E (20 mM HEPES/KOH pH 7.0, 2 mM Mg(OAc)_2_, 300 mM KOAc, 10% glycerol). Eluted protein was concentrated with 10 kDa cut-off Amicon ultra centrifugal filters before aliquoting, snap-freezing in liquid nitrogen and storage at −80°C.

#### PIAS4

The open reading frame of PIAS4 was codon optimized and cloned in a pNIC plasmid in frame with a *N*-terminal TwinStrep-ZB-tag. Recombinant PIAS4 protein was expressed in BL21 (DE3) *E. coli* cells, grown in TB (prepared with tap water) at 37°C to OD 0.7 before induction with 1 mM IPTG and expression at 18°C overnight. Protein purification was done analogously to SPRTN.

#### UBE2D3

For purification of UBE2D3, the open reading frame was codon optimized and cloned into a pDEST17 plasmid carrying an *N*-terminal His6-tag. UBE2D3 was expressed in BL21(DE3) *E. coli* cells, grown in TB media (prepared with tap water) to an OD of 0.7 at 37°C. Expression was induced by the addition of 0.5 mM IPTG for 3 h at 37°C. Cells were harvested, snap frozen in liquid nitrogen and stored at −80°C. All subsequent steps were performed at 4°C. For protein purification, cell pellets were thawed and resuspended in 50 mL buffer A (50 mM Na_2_HPO_4_/NaH_2_PO_4_ pH 8.0, 150 mM NaCl, 10 mM imidazole, 1 mM TCEP, cOmplete protease inhibitor, 0.04 mg/mL Pefabloc SC) and lysed by sonication. DNA was digested by the addition of smDNAse (45 U/mL lysate) for 30 min on a roller, followed by centrifugation at 18,000g for 30 min to remove cell debris. Clarified lysate was filtered using syringe filters (PVDF, 0.22 µm) and incubated with Ni-NTA agarose (QIAGEN) on a roller for 1 h at 4°C. The beads were washed with 20 mL buffer B (50 mM Na_2_HPO_4_/NaH_2_PO_4_ pH 8.0, 500 mM NaCl, 20 mM imidazole, 1 mM TCEP) and eluted in 5 mL buffer C (50 mM Na_2_HPO_4_/NaH_2_PO_4_ pH 8.0, 500 mM NaCl, 250 mM imidazole, 1 mM TCEP). The eluted protein was dialyzed against buffer D (20 mM Tris/HCl pH 7.5, 150 mM NaCl, 10% glycerol, 0.5 mM TCEP) overnight followed by SEC on a Superdex 200 Increase 10/300 GL column equilibrated in buffer D. Eluted protein was concentrated using 10 kDa cut-off Amicon ultra centrifugal filters before aliquoting, snap-freezing in liquid nitrogen and storage at −80°C.

#### RNF4

For purification of recombinant RNF4, the open reading frame was codon optimized and cloned in a pNIC plasmid in frame with a *N*-terminal TwinStrep-ZB-tag. RNF4 protein was expressed in BL21 (DE3) *E. coli* cells, grown in TB prepared with tap water and purified analogously to SPRTN. For SEC, buffer E (50 mM HEPES/KOH pH 7.2, 150 mM KCl, 10% glycerol, 1 mM TCEP, pH 7.2) was used.

#### SUMO2

For purification of recombinant SUMO2, the open reading frame was codon optimized and cloned in a pBAD plasmid carrying a *N*-terminal His6-tag. SUMO2 was expressed in BL21 (DE3) *E. coli* cells and grown in TB (prepared with tap water) at 37°C to OD 0.7 before induction with 0.02% L-arabinose and expression at 18°C overnight. Cells were harvested the next day, snap-frozen in liquid nitrogen and stored at −80°C. All subsequent steps were performed at 4°C. For protein purification, cell pellets were thawed, resuspended in buffer A (50 mM Na_2_HPO_4_/NaH_2_PO_4_ pH 7.5, 500 mM NaCl, 10% glycerol, 30 mM imidazole, 0.2% Triton-X-100, 1 mM TCEP, cOmplete protease inhibitor, 0.04 mg/mL Pefabloc SC, 1 mg/mL lysozyme, 45 U/mL smDNAse) and incubated on a roller for 30 min. Cell lysate was sonicated for 15 min before removal of cell debris by centrifugation at 18,000 g for 30 min. Clarified lysate was filtered using syringe filters (PVDF, 0.22 µm) and incubated with Ni-NTA agarose (QIAGEN) on a roller for 1 h at 4°C. The beads were transferred to a gravity flow column and washed with 15 CV buffer B (50 mM Na_2_HPO_4_/NaH_2_PO_4_ pH 7.5, 500 mM NaCl, 10% glycerol, 30 mM imidazole) before elution in 2 CV buffer C (50 mM Na_2_HPO_4_/NaH_2_PO_4_ pH 7.5, 500 mM NaCl, 10% glycerol, 300 mM imidazole). The His-tag was cleaved by the addition of His-tagged TEV protease (1 mg/L culture) and dialyzed against buffer D (20 mM HEPES/KOH pH 7.5, 100 mM KCl) overnight. The next day, cleaved protein was passed through Ni-NTA agarose to remove His-tagged TEV protease, residual uncleaved protein and the His-Tag. Flow-through was concentrated to 1 mL and loaded on a Superdex 200 Increase 10/300 GL column equilibrated in buffer E (20 mM HEPES/KOH pH 7.5, 100 mM KCl, 10% glycerol, 1 mM TCEP). Eluted protein was concentrated with 3 kDa cut-off Amicon ultra centrifugal filters before aliquoting, snap-freezing in liquid nitrogen and storage at −80°C.

### In vitro SPRTN autocleavage assays

Reactions were performed in 20 µL at 25°C for the indicated time, containing SPRTN (WT or variants, 250 nM), ΦX174 Virion ssDNA (11.14 nM) (NEB) and mono-ubiquitin (Ub^1^), Ub^1^-I44A (1 or 10 µM), M1-tetra-ubiquitin (Ub^4^), K48-Ub^4^, K63-Ub^4^, K63-Ub^3^ or K63-Ub^2^ (1 µM – referring to the concentration of individual ubiquitin moieties) (lifesensors). To examine *in trans* autocleavage assays, indicated Flag-SPRTN^FL^ variants (250 nM) were added additionally. The reaction buffer comprised 17.5 mM HEPES/KOH pH 7.2, 82.5 mM KCl, 3.25% glycerol and 5 mM TCEP. Reactions were stopped with 4x LDS sample buffer supplemented with 5% β-mercaptoethanol (β-ME), boiled at 95°C for 10 min and resolved on SDS-PAGE gels (4-12% Bis-Tris NuPAGE) using MOPS buffer and stained with SYPRO Ruby (Thermo Scientific) following manufacturer’s instructions. Gels were photographed using a BioRad Chemidoc MP system and cleavage was quantified using ImageJ (v1.54f), by dividing amounts of SPRTN by amounts of SPRTN present at the beginning of the reaction.

For experiments including Flag-SPRTN, cleavage of Flag-SPRTN was additionally evaluated by immunoblotting using anti-Flag antibody. Cleavage was quantified using ImageJ (v1.54f), by dividing amounts of Flag-SPRTN by amounts of Flag-SPRTN present at the beginning of the reaction.

### Flag pull down assays

Reactions were performed in 50 µL on ice for 15 min, containing indicated Flag-SPRTN variants (1 µM) and K63-Ub^4^ (0.5 µM, lifesensors). Ubiquitin concentrations refer to the concentration of individual ubiquitin moieties. The reaction buffer contained 5.7 mM HEPES/KOH pH 7.2,115 mM KCl, 3.1% glycerol, 0.5 mM TCEP and 0.1% IGEPAL. After incubation on ice, reactions were transferred to Anti-Flag® M2 magnetic beads (25 μL per sample) (Sigma) equilibrated with Flag-wash buffer (10 mM HEPES pH 7.2, 100 mM KCl, 2% glycerol, 0.1% IGEPAL). Reactions were incubated for 5 min on a rotating wheel at 4°C and washed 7 times with 500 μL Flag-wash buffer each. Bound protein was eluted with 30 µL 3xFLAG Peptide (100 ng/μL) (Sigma) by incubating for 30 min on a rotating wheel at 4°C. Supernatants were collected and boiled with 4x LDS sample buffer supplemented with 5% β-ME for 10 min at 95°C. Samples were resolved on SDS-PAGE gels (4-12% Bis-Tris NuPAGE) using MOPS buffer and subsequently immunoblotted using anti-SPRTN and anti-Ub antibodies.

### In vitro HMCES-DPC generation

DPCs were generated between HMCES^SRAP^ (WT or R98E), HMCES^SRAP^-K48-Ub^[short]/[long]^, HMCES^SRAP^-K63-Ub^[short]/[long]^ or HMCES^FL^ and a 30nt Cy5-labeled forward oligonucleotide (oDY_54), as previously described^44,45^. For HMCES^FL^-DPCs final concentrations differed from published protocols: HMCES^FL^ (13 μM), UDG (0.1 U/μL, NEB), DNA (1.25 μM). The crosslinking reaction was shortened to 30 min at 37°C. To form ssDNA-dsDNA junctions 1 µL complementary 15nt reverse oligonucleotide oHR_127 (12 µM in nuclease-free H_2_O) was annealed to all crosslinking reactions. For heat-denaturation (Figure S5C), DPCs were incubated for 5 min at 60°C.

### HMCES^SRAP^ Ubiquitylation using synthetic ubiquitin E3 ligases

A simplified Ubiquiton system^46^, based on fusions of a complete ubiquitin instead of split-ubiquitin as a starting point, was used. In brief, HMCES^SRAP^-Ub(G76V)-3C-FKBP-His6 was K48-poly-ubiquitylated in a reaction containing substrate (10 µM), ubiquitin (30 µM) (Merck KgaA), Ub^1^-K48R (10 µM), His-Uba1 (50 nM), Ubc7-His (E2) (4 µM), His-FRB-E3^48^ (10 µM), ATP (1 mM) and rapamycin (50 µM) in ubiquitylation buffer (40 mM HEPES/NaOH pH 7.4, 50 mM NaCl, 8 mM Mg(OAc)_2_) for 6.5 h at 30°C. K48-modified HMCES^SRAP^ was separated from other reaction components by cleaving the dimerization tag using His-3C-protease at 4°C overnight, reverse immobilized metal affinity chromatography (IMAC) and SEC (20 mM HEPES/KOH pH 7.8, 150 mM KCl, 2 mM MgCl_2_, 1 mM TCEP, 10% glycerol).

HMCES^SRAP^-Ub(G76V)-3C-FKBP-His6 was K63-poly-ubiquitylated in a reaction containing substrate (10 µM), ubiquitin (30 µM) (Merck KgaA), Ub^1^-K63R (10 µM), His-Uba1 (50 nM), His-Ubc13·Mms2 (E2) (2 µM), His-FRB-L20-E3^63^ (10 µM), ATP (1 mM) and rapamycin (50 µM) in ubiquitylation buffer (40 mM HEPES/NaOH pH 7.4, 50 mM NaCl, 8 mM Mg(OAc)_2_) for 2 h at 30°C and purified as described above.

Ubiquitin mutants, His-Uba1, Ubc7-His, His-Ubc13 and Mms2 were purified as previously described^54,55^. His-FRB-E3^48^ and His-FRB-L20-E3^63^ were produced in *E. coli* and purified by IMAC followed by SEC (20 mM HEPES/NaOH pH 7.4, 150 mM NaCl, 10% glycerol, 1 mM DTT) using an automated NGC chromatography system (Bio-Rad Laboratories).

### In vitro SUMOylation and ubiquitylation of HMCES-DPCs

SUMOylation reactions were performed in 20 µL for 30 min at 37°C, containing HMCES-DPC (125 nM), SUMO2 (1.250 µM), SAE1/UBA2 (100 nM) (Hölzel-Biotech), UBC9 (200 nM) and PIAS4 (125 nM). The reaction buffer comprised 20 mM HEPES/KOH pH 7.5, 110 mM KOAc, 5.32 mM MgCl2, 2 mM Mg(OAc)_2_, 0.05% TWEEN20, 0.2 mg/ml BSA, 1 mM TCEP, 2.5 mM ATP. If no further reactions were carried out 5 μL reaction buffer were added and DPCs were either used in DPC cleavage assays or directly mixed with 4x LDS sample buffer supplemented with 5% β-ME, followed by boiling for 1 min at 95°C prior to SDS-PAGE analysis. For subsequent ubiquitylation, 5 μL ubiquitin master mix were added, and reactions were incubated for 30 min at 37°C. The ubiquitin master mix contained ubiquitin (1 μM), UBE1 (100 nM) (lifesensors), RNF4 (200 nM) and UBE2D3 (200 nM). DPCs were either used in DPC cleavage assays or directly mixed with 4x LDS sample buffer supplemented with 5% β-ME, followed by boiling for 1 min at 95°C prior to SDS-PAGE analysis. Samples were resolved on SDS-PAGE gels (12% Bis-Tris BOLT) using MOPS buffer. Gels were scanned using a BioRad Chemidoc MP system with appropriate filter settings for Cy5 fluorescence. Gels were subsequently analyzed by immunoblotting using anti-K48-Ub and anti-K63-Ub antibodies.

For analysis of SUMOylated and ubiquitylated HMCES^FL^-DPC by mass spectrometry (MS), reactions were scaled up to 50 µL, ubiquitin concentration was increased (5 µM) and incubation time for ubiquitylation was extended (1 h at 37°C). Reactions were stopped by addition of 4x LDS sample buffer supplemented with 5% β-ME. Samples were stored at −20°C until mass spectrometry analysis.

### DPC cleavage assay

DPC cleavage by SPRTN was assessed in 10 µL reactions at 30°C for 1 h, containing SPRTN (WT or variants, as indicated – concentrations ranging from 0.1-100 nM), DPC or free DNA (10 nM) and optionally FANCJ (100 nM) and K63-Ub^4^ (0.4 or 1.6 µM, concentrations correspond to the concentration of individual ubiquitin moieties) (lifesensors). The reaction buffer comprised 17.1 mM HEPES/KOH pH 7.5, 85.6 mM KCl, 3.1% glycerol, 5 mM TCEP, 2.1 mM MgCl_2_, 0.12 mg/ml BSA and 1 mM ATP. Reactions were stopped with 4x LDS sample buffer supplemented with 5% β-ME and boiling for 1 min at 95°C. Samples were resolved on SDS-PAGE gels (12% Bis-Tris BOLT) using MOPS buffer. Gels were scanned using a BioRad Chemidoc MP system with appropriate filter settings for Cy5 fluorescence. DPC cleavage was quantified using ImageJ (v1.54f), by dividing the amount of cleaved DPCs by the total amount of DPC (cleaved + uncleaved). For Cleaved DPC*, the sum of cleavage fragment signals and the corresponding signal for free DNA was calculated minus free DNA signals inferred from control DPC reactions.

### Cellular SPRTN autocleavage assays

For cellular SPRTN autocleavage assays, 1 × 10^6^ cells were seeded in 6-well. 24 h later 4 µL siRNA (20 µM) and 20 µL Lipofectamine RNAiMAX Transfection Reagent (Thermo Scientific) were each diluted in 200 µL Opti-MEM serum-free medium. Following a 5 min incubation, siRNA and Lipofectamine RNAiMAX Transfection Reagent dilutions were mixed. After an additional 15 min, the transfection mix was added to cells. 24 h after transfection, each well was split in 4 wells of a 12-well plate. The next morning, cells were treated with 5-azadC (10 µM) (Sigma) for 2, 4 or 8 h or left untreated for each transfected siRNA. At desired time points, cells were directly lysed in 1x LDS and boiled for 20 min at 95°C. Samples were resolved on SDS-PAGE gels (4-12% Bis-Tris NuPAGE) using MOPS buffer and subsequently immunoblotted using anti-DNMT1, anti-SPRTN, anti-RNF4 and anti-Vinculin antibodies.

### Purification of x-linked proteins (PxP)

For PxP experiments, 7.5 × 10^5^ cells were seeded in 6-cm dishes, and thymidine-containing media (2 mM) (Sigma) was added after 8 h. After approximately 16 h, thymidine media was removed, and cells were washed twice with PBS and released into normal media for 9 h, before thymidine media was re-added and expression of YFP-SPRTN-Strep-tag variants was induced with DOX (1 µg/mL). After another 16 h in thymidine media, cells were washed twice with PBS and released into normal media for 2 h before adding fresh media containing 5-azadC (10 µM) (Sigma). After a 30 min incubation, 5-azadC containing media was removed, cells were washed twice with PBS and recovery was allowed for 2 h. Cells were scraped on ice at indicated time points and cell pellets were stored at −80°C. PxP to isolate DNMT1-DPCs was performed as previously described^10,48^. In brief, 10 µL of each cell suspension were directly lysed in 1x LDS sample buffer to serve as input samples before plug casting. 1.5 × 10^6^ cells were embedded into low-melt agarose (Bio-Rad) plugs, extracted by PxP^10,48^ and prepared for analysis by SDS-PAGE at the end of the protocol. Samples were resolved on SDS-PAGE gels (4-12% Bis-Tris NuPAGE) using MOPS buffer and subsequently immunoblotted using anti-DNMT1, anti-SPRTN, and anti-β-actin antibodies.

### Western blotting

Cells were lysed in NP-40 lysis buffer (50 mM Tris-HCl, pH 7.4, 150 mM NaCl, 0.1% NP-40 (IGEPAL), 10% glycerol, 5 mM EDTA, 50 mM NaF, 1 mM Na_3_VO_4_) supplemented with protease inhibitor cocktail (Sigma). Cell lysates containing 30 µg protein were resolved on SDS-PAGE gels (4-12% Bis-Tris NuPAGE) using MOPS buffer. Resolved proteins were transferred onto a nitrocellulose membrane, which was blocked for 1 h in 5% milk in Tris-buffered saline supplemented with 0.1% Tween-20 (TBS-T) and analyzed by standard immunoblotting using antibodies listed in key resources.

### Cell viability

For cell proliferation assays, 1,000 cells were seeded per well (n=8) in a 96-well plate, and cell numbers were recorded every 8 h for 3 days using Cytation 5 (BioTek) equipped with a 4x objective and the Gen5 software (ver. 3.14).

### Flow cytometry

Cells were labeled with EdU (10 μM) for 45 min. EdU staining was performed with the Click-iT EdU Alexa Fluor 488 Flow Cytometry Assay Kit (Thermo Scientific) following the manufacturer’s protocol. Cells were next stained with 4’,6-diamidino-2-phenylindole (DAPI) (4 μg/mL) (Thermo Scientific) and analyzed using the BD LSRFortessa cell analyzer (BD Biosciences) with the FACSDiva^TM^ software (ver. 6.2). Figures were generated using FlowJo (ver. 10.10).

### Microscopy

Cells grown on a cover glass were washed once with PBS, fixed with paraformaldehyde (4%) in PBS for 10 min, and permeabilized with 0.2% Triton X-100 in PBS for 10 min. After washing with PBS, cells were stained with DAPI (1 μg/mL) (Thermo Scientific) in PBS for 10 min, and the cover glass was mounted with ProLong Glass Antifade Mountant (Thermo Scientific). Images were captured using a Zeiss Axio Observer 7 equipped with Apotome 3 and the Axiocam 820 camera. At least 300 DAPI-stained nuclei were scored manually for the presence of micronuclei or chromatin bridges by an observer blinded to sample identities. Statistical analysis was performed by two-way ANOVA with Dunnett’s multiple comparison test in GraphPad Prism (ver. 10.3.0).

### Identification of ubiquitin-linkages by quantitative mass spectrometry analysis

*In vitro* DPC ubiquitylation reactions were terminated by boiling samples at 70°C. For quantitative mass spectrometry analysis, samples were subsequently cleaned-up using the paramagnetic-based SP3 technology as described previously^56^. Briefly, 100 µg of freshly pre-equilibrated SP3 beads were added to 20 µL of samples. Purification of total proteins from *in vitro* reactions was next completed through three-rounds of 80% (v/v) ethanol-solvation of the SP3-sample mixture at room temperature. The resulting purified proteins were then subjected to trypsin digestion (1 µg) in 50 mM ammonium bicarbonate pH 8.0 for 16 h at 37°C. Digested peptides were acidified using trifluoroacetic acid and desalted on reverse-phase C18 StageTips for MS analysis. Eluted samples were analyzed on a quadrupole Orbitrap mass spectrometer Exploris 480 (Thermo Scientific) equipped with a UHPLC EASY-nLC 1200 system (Thermo Scientific). Samples were loaded onto a C18 reversed-phase column (55 cm length, 75 mm inner diameter, packed in-house with ReproSil-Pur 120 C18-AQ 1.9-µm beads, Dr. Maisch GmbH) and eluted with a gradient from 2.4 to 32% Acetonitrile.

The mass spectrometer was operated in data-dependent mode, automatically switching between MS and MS2 acquisition. Survey full scan MS spectra (m/z 300–1,650; resolution: 60,000; target value: 3 × 10^6^; maximum injection time: 28 ms) were acquired in the Orbitrap. The 15 most intense precursor ions were sequentially isolated, fragmented by higher energy C-trap dissociation (HCD) and scanned in the Orbitrap mass analyzer (normalized collision energy: 30%; resolution: 15,000; target value: 1 × 10^5^; maximum injection time: 40 ms; isolation window: 1.4 m/z (LFQ run)). Precursor ions with unassigned charge states, as well as with charge states of +1 or higher than +6, were excluded from fragmentation. Precursor ions already selected for fragmentation were dynamically excluded for 25 s.

Peptide identification: Raw data files were analyzed using MaxQuant (development version 1.5.2.8). Parent ion and MS2 spectra were searched against a database containing 98,566 human protein sequences obtained from UniProtKB (April 2018 release) using the Andromeda search engine. Spectra were searched with a mass tolerance of 6 ppm in MS mode, 20 ppm in HCD MS2 mode and strict trypsin specificity, allowing up to three miscleavages. Protein N-terminal acetylation and methionine oxidation were searched as variable modifications. The dataset was filtered based on posterior error probability (PEP) to arrive at a false discovery rate (FDR) of less than 1% estimated using a target-decoy approach.

### ICP-OES measurements

SPRTN samples in storage buffer and expression media (TB) as control were digested using an *Anton Paar* Multiwave 5000 microwave. For this, 160 µL of each sample (corresponding to 0.95 mg protein) were placed into PTFE digestion vessels. To this, 1 mL HNO_3_ (69%) (VWR) was added. The used digestion program was: 5 min ramp up to 180°C, then 10 min at 200°C and 15 min at 220°C. After digestion, samples were allowed to cool to room temperature before being diluted to final volumes (10 mL) with ultrapure water (type 1, 18.2 MΩ·cm at 25°C) for Inductively Coupled Plasma Optical Emission Spectrometry (ICP-OES) analysis. Blank samples were treated analogously.

ICP-OES was performed on a Varian Vista RL instrument operating in radial mode to determine the concentrations of Co, Fe, Mn and Zn. Calibration standards were prepared in HNO_3_ (2%) by diluting a certified multi-element ICP standard containing the elements of interest (Merck) to obtain a 4-point linear calibration curve ranging from 0 µg/mL to 4 µg/mL. ICP-OES operating conditions were set as follows: plasma power at 1.25 kW, plasma gas at 13.5 L/min, nebulizer pressure at 170 kPa, auxiliary gas flow rate at 1.5 L/min with three replicates per measurement cycle, which were automatically averaged. The following emission lines were selected for Co at 230.786, 231.160, 237.863 and 258,033 nm, Fe at 234.350, 238.204, 258.588 and 259.940 nm, Mn at 257.610, 259.372 and 293.931 nm and Zn at 202.548, 206.200 and 213.857 nm. Quality control was ensured by analyzing blanks within the sequence and a certified reference material alongside the samples, with recoveries within acceptable limits.

### Protein structure predictions

To prepare the crystal structure (PDB: 6mdx) for MD simulations, Swiss Model^57^ was employed to model the two missing residues, to adjust the modified amino acids to their natural counterpart and to remove the ligands from crystallization. From the structures generated with Swiss Model, we took the one that was the closest to the crystal structure with an RMSD of 0.082 Å.

Structures for SprT-BR (SPRTN^aa28–245^) and SprT-BR-ubiquitin were generated using ColabFold^43,58,59^ and AlphaFold2^60^ using alphafold2_ptm. Figures were generated using PyMOL (ver. 3.0.3).

### Molecular dynamics simulations

Starting structures for SprT and SprT-Ubiquitin were generated as described above, in both cases the top ranked model (Rank_1) was selected for further analysis. Predictions for SPRTN variants (L38S, L99S) were generated analogously. In case of SprT-ubiquitin complexes, the interface predicted for the WT enzyme was also used for SPRTN variants. Starting with these predicted structures, two Zn^2+^ ions were added based on their binding sites in the crystal structure. Subsequently, hydrogen atoms were added to these structures as well as to the model of the crystal structure employing the H++ server^61^. This way, the model for SprT and for the crystal structure consisted of the same atoms and of amino acids in the same protonation states. Figures were generated using PyMOL (ver. 3.0.3).

For setting-up the system for the simulations, we employed AmberTools20^62^. The structures were placed in rectangular simulation boxes with a minimum distance of 15 Å between the solute and the boundaries of the box. The boxes were filled with water and NaCl was added to neutralize the system and to achieve a physiological concentration of around 150 mM NaCl leading to system sizes of about 72.324 to 95.881 atoms for SprT and the SprT-ubiquitin complex, respectively.

The force field parameters for the proteins were taken from the Amber ff19SB force field^63^ and the OPC water model was used^64^. After conversion of the topology and coordinate files to gromacs with parmed from AmberTools20, the parameters for NaCl were replaced by the ones from Ref^65^ and for Zn^2+^ by the parameters from Ref^66^.

MD simulations were performed using the Gromacs simulation package^67^, version 2024. Initially, the energy of the systems was minimized using the steepest descent algorithm. Subsequently, the systems were equilibrated for 1 ns, first in the NVT and then in the NPT ensemble. For the production run, we performed 400 ns long simulations employing the velocity-rescaling thermostat with a stochastic term and a time constant of 0.1 ps and isotropic Parrinello-Rahman pressure coupling with a time constant of 5.0 ps. Each production was repeated three times with random velocities. We used clustering analysis with Gromacs for the production runs based on RMSD to group similar conformations, allowing us to identify the dominant structural states and to calculate the radius of gyration (Rg) of the clusters. In this analysis, 100 ns were discarded for equilibration.

To calculate the binding affinity, additional simulations were performed. First 1 ns of NVT and NPT simulations with stronger restraints of 2.1 kcal/mol Å^2^. Then production runs with weak restraints of 0.1 kcal/mol. We used representative structures from the largest cluster of the unrestrained simulations as starting structures. The productions were performed for 4 ns neglecting the first ns for equilibration and structures were sampled every 10 ps. Each simulation was repeated three times. The end-point free energy calculations were performed using the MMPBSA program from the Amber package^68^ using the gmx_MMPBSA tool^69^. Water molecules and ions were removed and the trajectories were re-evaluated using the mbondi3 radii and parameters from Ref^70^ denoted as igb=8 in Amber. To account for hydrophobic solvation, we used a surface area-dependent tension model with surface tension coefficient γ=0.005 kcal/mol Å^2^. No conformational changes were considered. From the alanine scanning procedure, we obtained the contribution of the two mutations to the binding free energy. The energy contribution of the mutation was calculated by cutting all atoms after the Cβ-atom of this residue. This procedure was performed for the simulations of the WT and the simulations of the mutated complexes. The difference yielded the binding energy contributions of the L38S and the L99S mutations.

### NMR spectroscopy

All the NMR samples (uniformly labeled ^13^C,^15^N-SPRTN^151–245, 15^N-SPRTN^28–245^-E112Q ± L99S, ^15^N-SPRTN^151–214^) were prepared at concentrations of 100 µM and 250 µM in 20mM HEPES/KOH pH 7.2, 150 mM KCl, 2 mM TCEP with 10% D_2_O, as lock signal. All NMR experiments were recorded at 308 K on a 600 MHz Bruker Avance NMR spectrometer, equipped with a cryogenic triple resonance gradient probe. NMR spectra were processed using NMRPIPE^71^ or TOPSPIN3.7 (Bruker) and analyzed using NMRFAM-SPARKY^72^. Using the previous backbone resonance assignments for SPRTN^151–245^ from Ref^32^, aromatic resonances were further assigned using 2D CBHD, CBHE, aromatic ^1^H,^13^C-HSQC and 3D ^15^N/^13^C-edited NOESY experiments. For 2D ^1^H,^15^N-HSQC comparisons, 100 µM SPRTN was mixed with 500 µM Ub^1^ (5x molar excess) / 250 µM Ub^1^-I44A (2.5x molar excess) and / or 200 µM dsDNA (2x molar excess), accordingly. Chemical shift perturbations (CSP) values were calculated based as 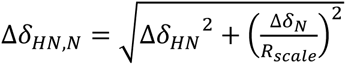, where *R*_scale_=6.5 was applied as suggested previously^73^.

## ACKNOWLEDGMENTS

We thank B. Schulman for providing a plasmid for recombinant mono-ubiquitin. S.D. and P.W. are supported by the International Max-Planck Research School for Molecules of Life. We are grateful to Sam Asami and Gerd Gemmecker for the help with NMR measurements at the Bavarian NMR center. We thank Dr. Shar-yin N. Huang at the National Cancer Institute for her technical support. Research in the lab of J.S. is funded by European Research Council (ERC StG 801750 DNAProteinCrosslinks, ERC CoG 101124695 DECONSTRUCT), the Alfried-Krupp von Bohlen und Halbach-Stiftung, European Molecular Biology Organization (YIP4644), a Vallee Foundation Scholarship, and Deutsche Forschungsgemeinschaft (Project ID 213249687 - SFB 1064). H.D.U. acknowledges funding by the European Research Council (ERC AdG 101140447). We acknowledge funding by the Deutsche Forschungsgemeinschaft (J.S., H.D.U., and P.B.: Project-ID 393547839 – SFB 1361; M.S. and L.J.D: Project-ID 325871075 – SFB1309). The authors acknowledge the scientific support and HPC resources provided by the Erlangen National High Performance Computing Center (NHR@FAU) of the Friedrich-Alexander-Universität Erlangen-Nürnberg (FAU) under the NHR project b119ee and the resources on the LiCCA HPC cluster of the University of Augsburg, co-funded by the Deutsche Forschungsgemeinschaft under Project-ID 499211671).

## AUTHORS CONTRIBUTIONS

**Sophie Dürauer:** Conceptualization; Data curation; Formal analysis; Investigation; Methodology; Validation; Visualization; Writing – original draft; Writing – review & editing. **Hyun-Seo Kang:** Data curation; Formal analysis; Investigation; Methodology; Validation; Visualization; Writing – review & editing. **Christian Wiebeler:** Data curation; Formal analysis; Investigation; Methodology; Software; Validation; Visualization; Writing – review & editing. **Yuka Machida:** Investigation; Validation; Writing – review & editing. **Dina S Schnapka:** Investigation; Methodology; Validation; Writing – review & editing. **Denitsa Yaneva:** Investigation; Validation; Writing – review & editing. **Christian Renz:** Investigation; Methodology; Writing – review & editing. **Maximilian J Götz:** Methodology; Writing – review & editing. **Pedro Weickert:** Investigation; Validation; Writing – review & editing. **Aligail Major:** Formal analysis; Investigation; Methodology; Software; Validation. **Aldwin Suryo Rah:** Data curation; Formal analysis; Investigation; Writing – review & editing. **Sophie M Gutenthaler-Tietze:** Formal analysis; Investigation; Writing – review & editing. **Lena J Daumann:** Funding acquisition; Resources; Supervision; Writing – review & editing. **Petra Beli:** Funding acquisition; Resources; Supervision; Writing – review & editing. **Helle D. Ulrich:** Funding acquisition; Resources; Supervision; Writing – review & editing. **Michael Sattler:** Funding acquisition; Resources; Supervision; Writing – review & editing. **Yuichi J. Machida:** Formal analysis, Funding acquisition, Investigation; Resources; Supervision; Validation; Writing – review & editing. **Nadine Schwierz:** Formal analysis, Funding acquisition; Investigation; Methodology; Resources; Software; Supervision; Validation; Writing – review & editing. **Julian Stingele:** Conceptualization; Funding acquisition; Project administration; Resources; Supervision; Writing – original draft; Writing – review & editing.

---

**Figure S1.**
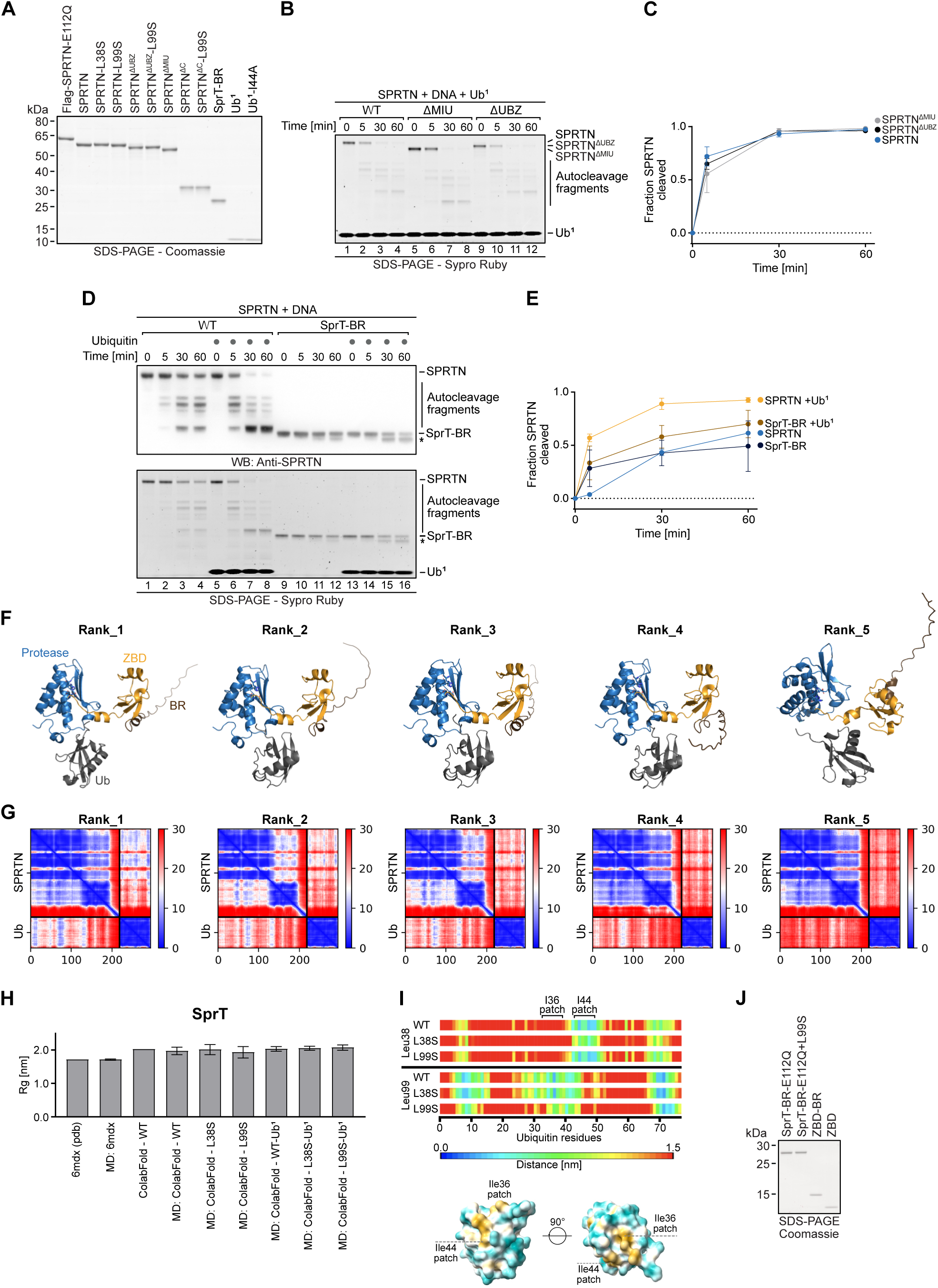
Ubiquitin boosts SPRTN’s protease activity independent of its established ubiquitin binding domains. (A) Coomassie stained SDS-PAGE gel, showing equimolar amounts of purified recombinant human Flag-SPRTN-E112Q, SPRTN, SPRTN-L38S, SPRTN-L99S, SPRTN^ΔUBZ^, SPRTN^ΔUBZ^-L99S, SPRTN^ΔMIU^, SPRTN^ΔC^, SPRTN^ΔC^-L99S, SprT-BR, mono-ubiquitin (Ub^1^) and Ub^1^-I44A used for *in vitro* assays. (B) Indicated recombinant SPRTN variants (250 nM) were incubated in the presence of Virion DNA (11.14 nM) and Ub^1^ (10 µM). (C) Quantification of SPRTN autocleavage assay shown in (B), depicting the mean ± SD of three independent experiments. (D) Indicated SPRTN variants (250 nM) were incubated alone or in the presence of Virion DNA (11.14 nM) with or without Ub^1^ (10 µM). SprT-BR autocleavage fragment is marked with asterisks. (E) Quantification of SPRTN autocleavage assay shown in (D), depicting the mean ± SD of three independent experiments. (F) Structures of SprT-BR-ubiquitin complex predicted by ColabFold using AlphaFold2_ptm. All five models (Rank_1-5) are shown (left to right), highlighting SPRTN’s protease domain (blue), Zinc-binding domain (ZBD) (orange) and basic region (BR) (brown). Ubiquitin (Ub) is shown in grey. (G) Predicted aligned error (PAE) blots of predicted SprT-BR-ubiquitin complexes shown in (F). (H) Bar charts showing Radius of gyration (Rg) before and after molecular dynamics (MD) simulations for SprT (PDB: 6mdx and ColabFold predicted) and SprT-Ub^1^ (ColabFold predicted). In case of MD simulations, the mean ± SD of snapshots from three independent 400 ns MD trajectories is given and the first 100 ns of each trajectory were discarded for equilibration. (I) Heat map indicating minimum distances of all amino acids in ubiquitin to residues at positions 38 (upper part) and 99 (lower part) of SPRTN, including wild-type (WT) conditions (L38, L99) and mutated states (L38S, L99S) (top). Structure of ubiquitin colored by hydrophobicity (bottom). Ile36 and Ile44 patches are highlighted. (J) Coomassie stained SDS-PAGE gel, showing equimolar amounts of purified recombinant human SprT-BR-E112Q, SprT-BR -E112Q+L99S, ZBD-BR, and ZBD used for NMR measurements.

**Figure S2.**
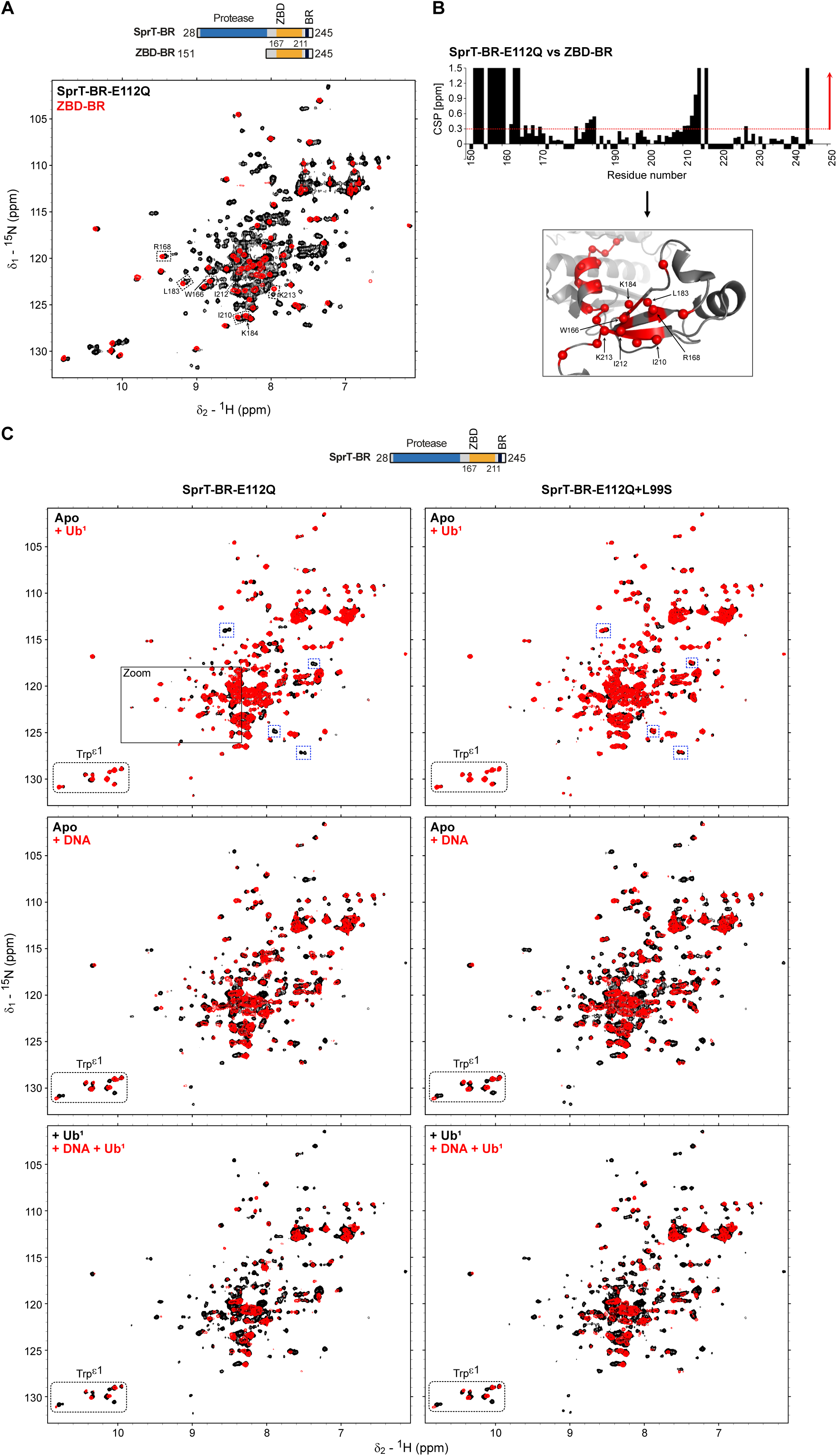
Ubiquitin binds to the SprT-domain via the USD. (A) Comparison of NMR spectral of catalytically inactive SprT-BR-E112Q (black) and SPRTN^151–245^ (red). Some peaks with Chemical Shift Perturbation (CSP) > 0.3 ppm are annotated and boxed. (B) CSP of SprT-BR-E112Q against ZBD-BR. Residues with CSP > 0.3 ppm (red arrow) are highlighted in the structure as red spheres with labels for some representative residues (bottom). Negative values indicate proline or unassigned residues. Large changes, which could not be traced are given a CSP value of 1.5 ppm. (C) Comparison of NMR spectra (Trp ε1 amides in ^1^H,^15^N-HSQC experiments) of SprT-BR-E112Q (left) and SprT-BR-E112Q+L99S (right), alone (black), with mono-ubiquitin (Ub^1^) (5x molar excess) (red) (top), with dsDNA (2x molar excess) (red) (middle) and of Ub^1^-bound (5x molar excess) SprT-BR-E112Q alone (black) and in the presence of dsDNA (5x molar excess, (red) (bottom). Trp ε1 region is labeled and shown with a box. Selective peaks are boxed in blue to highlight the spectral differences between SprT-BR-E112Q and SprT-BR-E112Q+L99S upon adding Ub^1^ (top). Zoom-In region in Figure S3B is marked with a black box (left) (top).

**Figure S3.**
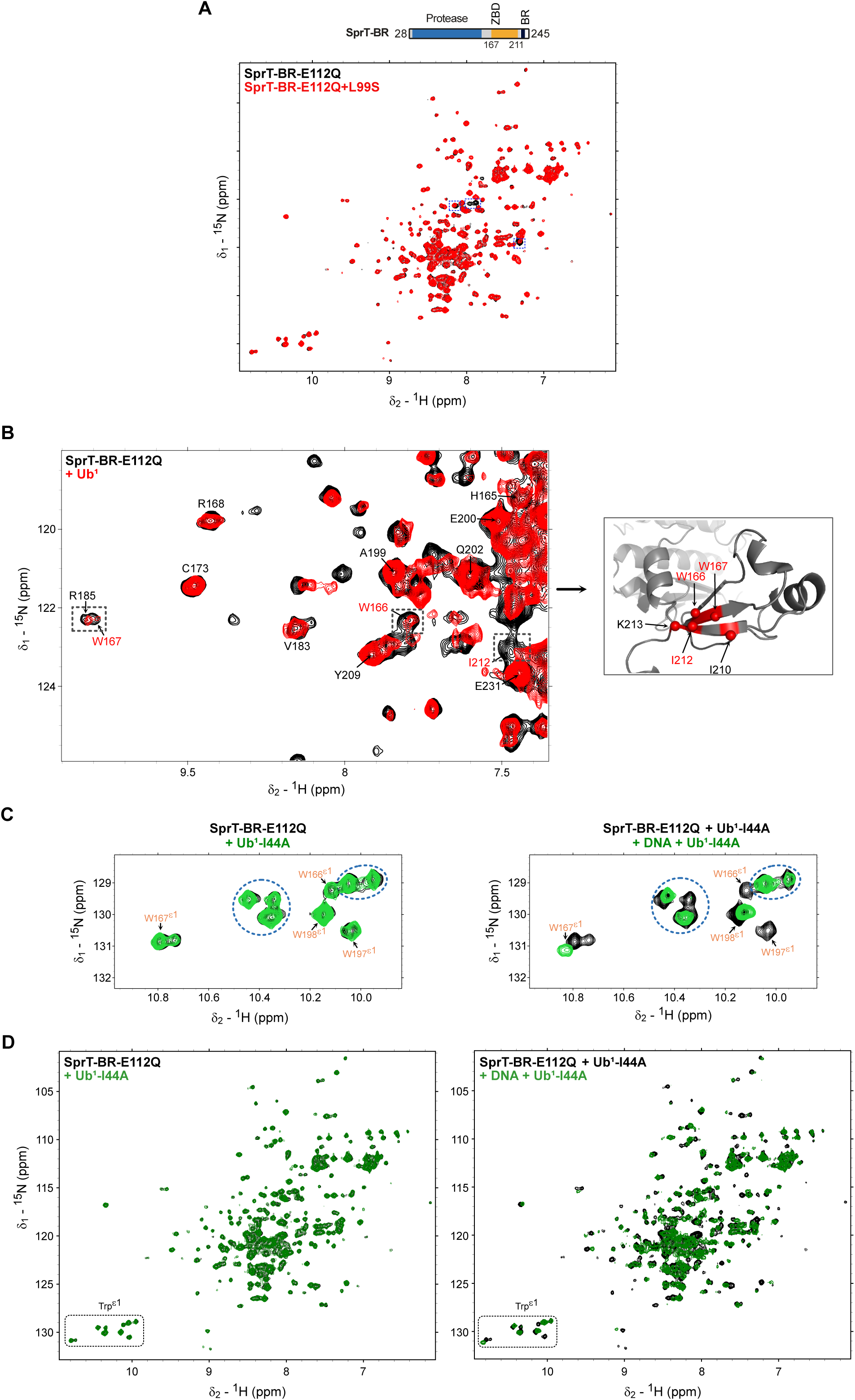
Ubiquitin binding results in conformational changes at the SprT domain. (A) Comparison of NMR spectra of SprT-BR-E112Q (black) and SprT-BR-E112Q+L99S (red). Spectral differences are highlighted with boxes. (B) Comparison of NMR spectra of SprT-BR-E112Q alone (black) and with mono-ubiquitin (Ub^1^) (5x molar excess) (red) (left). Resonances corresponding to the zinc-binding domain (ZBD) are labeled. The broadened ZBD signals in the presence of Ub^1^ are labeled in red with squares in the spectra (left) and highlighted in the structure (right). Full spectrum shown in Figure S2C (left) (top). (C) Comparison of NMR spectra (Trp ε1) of catalytically inactive SprT-BR-E112Q alone (black), with mono-ubiquitin (Ub^1^)-I44A (2.5x molar excess) (green) (left) and of Ub^1^-I44A-bound (2.5x molar excess) SprT-BR-E112Q alone (black) and in the presence of dsDNA (2x molar excess) (green) (right). Resonances corresponding to the Trp ε1’s in the ZBD are labeled in orange or shown as asterisk when broadened. Trp ε1’s in the protease domain are shown with blue circles. Full spectra are shown in panel (D). (D) Full NMR spectra of panel (C). Trp ε1 region is shown with a box.

**Figure S4.**
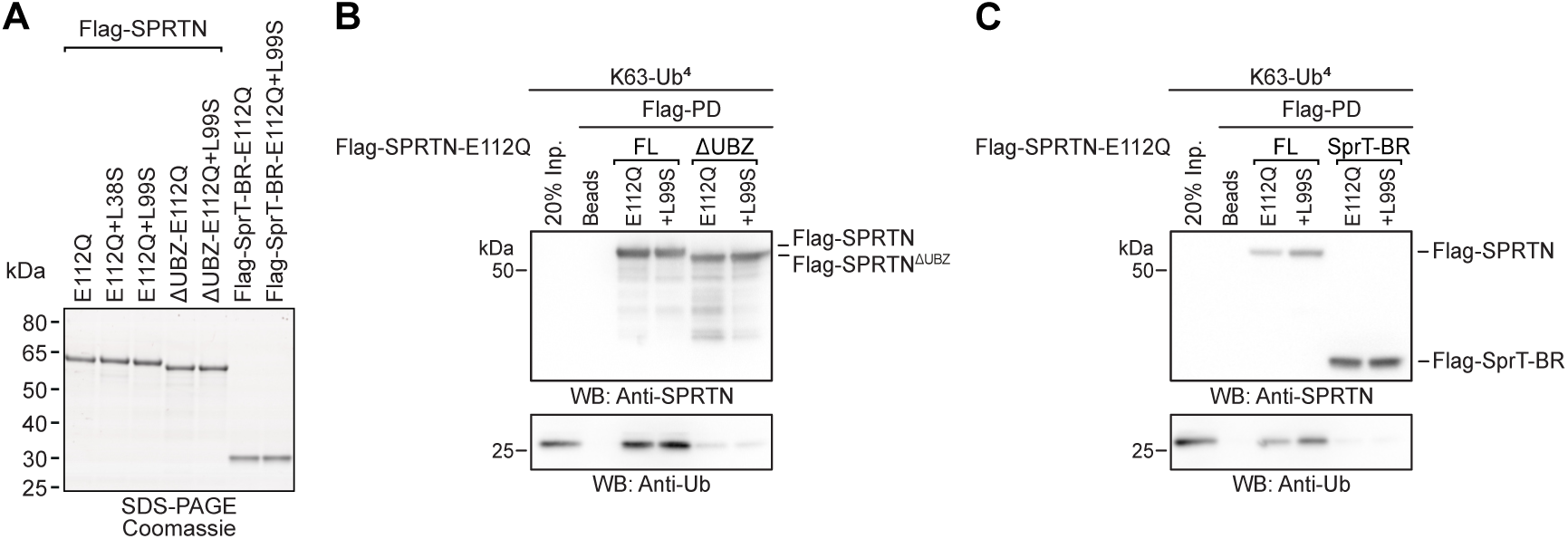
SPRTN’s UBZ mediates ubiquitin binding. (A) Coomassie stained SDS-PAGE gel, showing equimolar amounts of purified recombinant human Flag-SPRTN^FL^-E112Q, Flag-SPRTN^FL^-E112Q+L38S, Flag-SPRTN^FL^-E112Q+L99S, Flag-SPRTN^ΔUBZ^-E112Q, Flag-SPRTN^ΔUBZ^-E112Q+L99S, Flag-SprT-BR-E112Q, Flag-SprT-BR-E112Q+L99S used for *in vitro* assays. (B-C) Indicated recombinant Flag-SPRTN variants (1 µM) were incubated with K63-tetra-ubiquitin (Ub^4^) (0.5 µM) for 15 min on ice. Ubiquitin concentrations correspond to the concentration of individual ubiquitin moieties. Flag-SPRTN variants were pull downed (PD) together with bound ubiquitin using Anti-Flag beads. Reactions were analyzed by immunoblotting.

**Figure S5.**
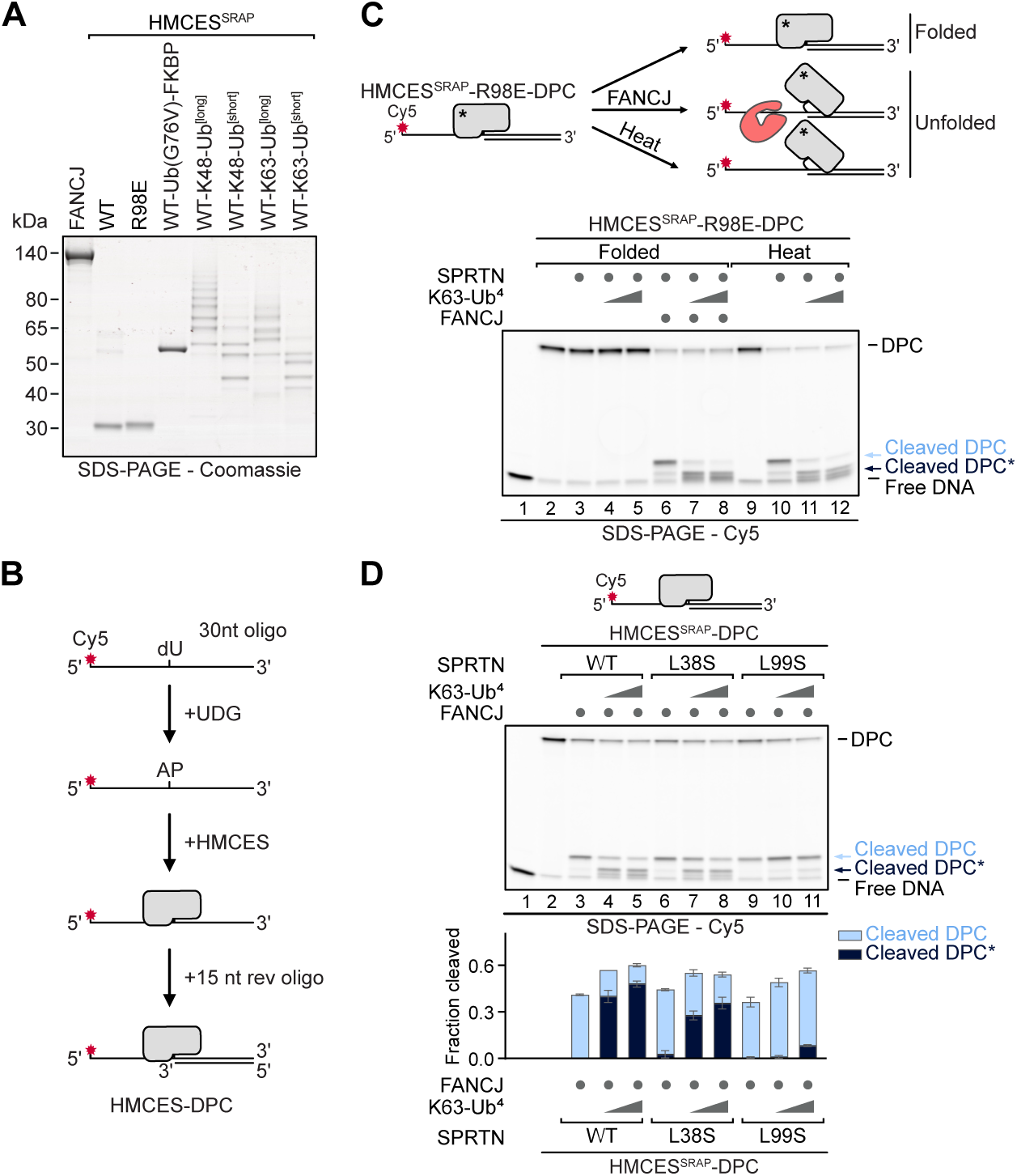
DPC ubiquitylation promotes cleavage by SPRTN. (A) Coomassie stained SDS-PAGE gel, showing equimolar amounts of purified recombinant human FANCJ, HMCES^SRAP^, HMCES^SRAP^-R98E, HMCES^SRAP^-Ub(G76V)-FKBP, HMCES^SRAP^-K48-Ub^[long]^, HMCES^SRAP^-K48-Ub^[short]^, HMCES^SRAP^-K63-Ub^[long]^ and HMCES^SRAP^-K63-Ub^[short]^ used for *in vitro* assays. (B) Schematic of the generation of HMCES-DPCs. HMCES was incubated for 30 min at 37°C with a Cy5-labeled 30nt oligonucleotide containing a dU at position 15 and UDG. After crosslinking a complementary 15nt reverse oligonucleotide was annealed to form a ssDNA-dsDNA junction. (C) HMCES^SRAP^-R98E-DPCs were either used in their native state (folded) or unfolded by addition of FANCJ or heat denaturation by incubating for 5 min at 60°C (top). Indicated HMCES^SRAP^-R98E-DPCs (10 nM) were incubated alone or in the presence of FANCJ (100 nM), SPRTN (100 nM) or indicated concentrations of K63-Ub^4^ (0.4 or 1.6 µM, referring to the concentration of individual ubiquitin moieties) for 1 h at 30°C. (D) Indicated HMCES^SRAP^-DPCs (10 nM) were incubated alone or in the presence of FANCJ (100 nM) and indicated variants of SPRTN (100 nM), with or without indicated concentrations of K63-Ub^4^ (0.4 or 1.6 µM, referring to the concentration of individual ubiquitin moieties) for 1 h at 30°C. Quantification: bar graphs represent the mean ± SD of three independent experiments.

**Figure S6.**
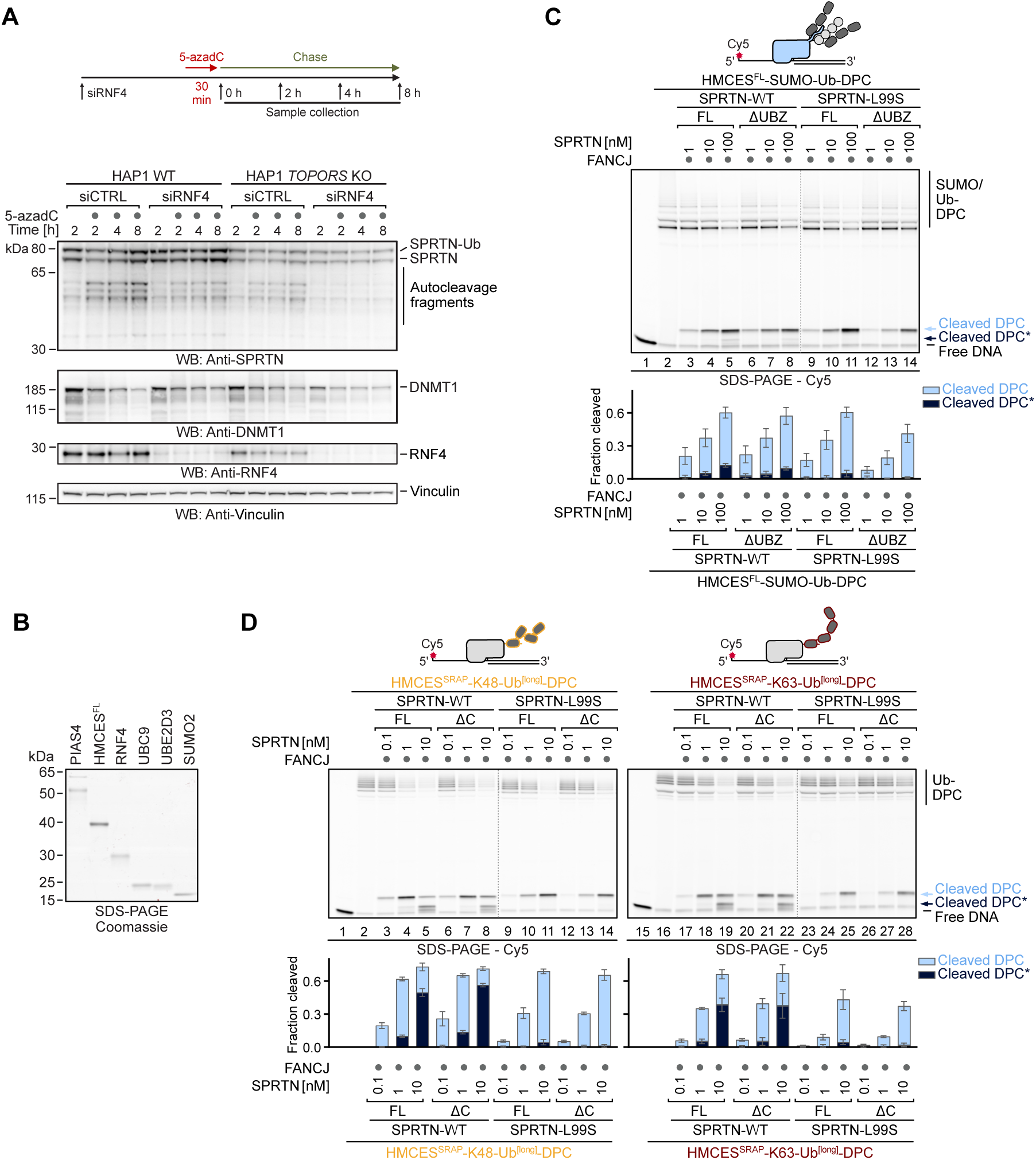
Global-genome DPC repair by SPRTN is mediated by the USD. (A) HAP1 wild-type (WT) or HAP1 *TOPORS* knock-out (KO) cells transfected with indicated siRNAs were treated with 5-azadC (10 µM) and harvested as depicted (top). Whole cell lysates were analyzed by immunoblotting (bottom). (B) Coomassie stained SDS-PAGE gel, showing equimolar amounts of purified recombinant human PIAS4, HMCES^FL^, RNF4, UBC9, UBE2D3 and SUMO2 used for *in vitro* assays. (C) HMCES^FL^-SUMO-Ub-DPCs (10 nM) were incubated alone or in the presence of FANCJ (100 nM) and indicated variants and concentrations of SPRTN^FL^ or SPRTN^ΔUBZ^ (1-100 nM) for 1 h at 30°C. Quantification of DPC cleavage assay: bar graphs represent the mean ± SD of three independent experiments. (D) Indicated HMCES^SRAP^-Ub^[long]^-DPCs (10 nM) were incubated alone or in the presence of recombinant FANCJ (100 nM) and indicated variants and concentrations of recombinant SPRTN^FL^ or SPRTN^ΔC^ (0.1-10 nM) for 1 h at 30°C. Quantification: bar graphs represent the mean ± SD of three independent experiments. Values for SPRTN^FL^ were also used for quantification of Figure 5G.

**Figure S7.**
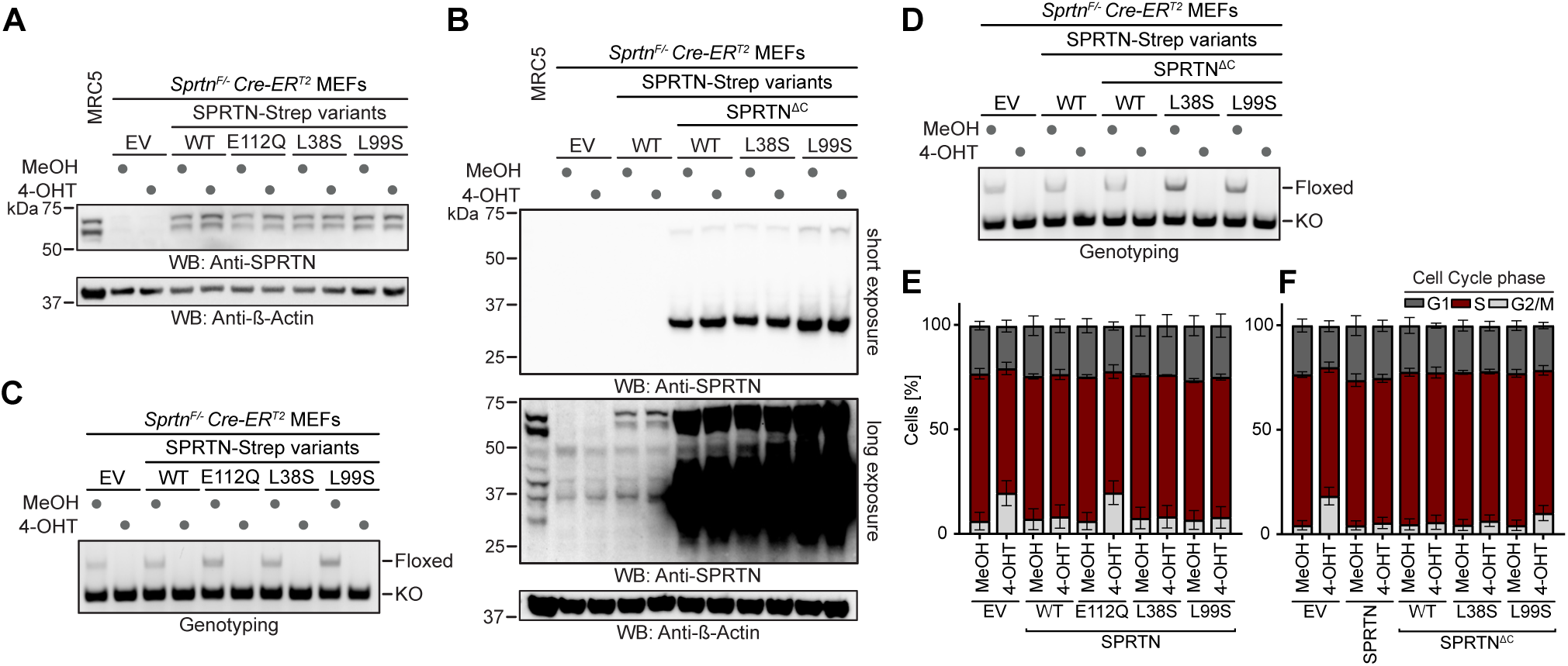
Ubiquitin-dependent activation of SPRTN maintains genome stability in Ruijs-Aalfs syndrome. (A-B) Expression of indicated SPRTN variants or empty vector (EV) (pMSCV) in *Sprtn^F/-^* Cre-ER^T2^ mouse embryonic fibroblasts (MEFs) treated with methanol (MeOH) or (Z)-4-hydroxytamoxifen (4-OHT) (2 µM) for 48 h. Whole cell lysates were anlayzed by immunoblotting. (C-D) PCR-based genotyping of Sprtn alleles in *Sprtn^F/-^* Cre-ER^T2^ MEFs, complemented with indicated SPRTN variants or EV (pMSCV), treated with MeOH or 4-OHT (2 µM) for 48 h. (E-F) Cell cycle profiling of *Sprtn^F/-^* Cre-ER^T2^ MEFs, complemented with indicated SPRTN variants or EV (pMSCV), treated with MeOH or 4-OHT (2 µM) for 48 h. Cells were labeled with EdU for 45 min and analyzed by flow cytometry. Bar charts represent the mean ± SD of three independent experiments.

## REFERENCES

1. Tubbs, A. & Nussenzweig, A. Endogenous DNA Damage as a Source of Genomic Instability in Cancer. Cell 168, 644–656 (2017).

2. Schumacher, B., Pothof, J., Vijg, J. & Hoeijmakers, J. H. J. The central role of DNA damage in the ageing process. Nature 592, 695–703 (2021).

3. Dantuma, N. P. & van Attikum, H. Spatiotemporal regulation of posttranslational modifications in the DNA damage response. EMBO J 35, 6–23 (2016).

4. Blackford, A. N. & Jackson, S. P. ATM, ATR, and DNA-PK: The Trinity at the Heart of the DNA Damage Response. Mol Cell 66, 801–817 (2017).

5. Weickert, P. & Stingele, J. DNA-Protein Crosslinks and Their Resolution. Annu Rev Biochem 91, 157–181 (2022).

6. Stingele, J., Bellelli, R. & Boulton, S. J. Mechanisms of DNA-protein crosslink repair. Nat Rev Mol Cell Biol 18, 563–573 (2017).

7. Wojtaszek, J. L. & Williams, R. S. From the TOP: Formation, recognition and resolution of topoisomerase DNA protein crosslinks. DNA Repair (Amst) 142, 103751 (2024).

8. Mehta, K. P. M., Lovejoy, C. A., Zhao, R., Heintzman, D. R. & Cortez, D. HMCES Maintains Replication Fork Progression and Prevents Double-Strand Breaks in Response to APOBEC Deamination and Abasic Site Formation. Cell Rep 31, 107705 (2020).

9. Tretyakova, N. Y., Groehler, A. & Ji, S. DNA-Protein Cross-Links: Formation, Structural Identities, and Biological Outcomes. Acc Chem Res 48, 1631–1644 (2015).

10. Weickert, P. et al. SPRTN patient variants cause global-genome DNA-protein crosslink repair defects. Nat Commun 14, 352 (2023).

11. Duxin, J. P., Dewar, J. M., Yardimci, H. & Walter, J. C. Repair of a DNA-protein crosslink by replication-coupled proteolysis. Cell 159, 346–357 (2014).

12. Carnie, C. J. et al. Transcription-coupled repair of DNA-protein cross-links depends on CSA and CSB. Nat Cell Biol 26, 797–810 (2024).

13. van Sluis, M. et al. Transcription-coupled DNA-protein crosslink repair by CSB and CRL4CSA-mediated degradation. Nat Cell Biol 26, 770–783 (2024).

14. Oka, Y., Nakazawa, Y., Shimada, M. & Ogi, T. Endogenous aldehyde-induced DNA-protein crosslinks are resolved by transcription-coupled repair. Nat Cell Biol 26, 784–796 (2024).

15. Liu, J. C. Y. et al. Mechanism and function of DNA replication-independent DNA-protein crosslink repair via the SUMO-RNF4 pathway. EMBO J 40, e107413 (2021).

16. Sun, Y. et al. A conserved SUMO pathway repairs topoisomerase DNA-protein cross-links by engaging ubiquitin-mediated proteasomal degradation. Sci Adv 6, eaba6290 (2020).

17. Carnie, C. J. et al. Decitabine cytotoxicity is promoted by dCMP deaminase DCTD and mitigated by SUMO-dependent E3 ligase TOPORS. EMBO J 43, 2397–2423 (2024).

18. Liu, J. C. Y. et al. Concerted SUMO-targeted ubiquitin ligase activities of TOPORS and RNF4 are essential for stress management and cell proliferation. Nat Struct Mol Biol 31, 1355–1367 (2024).

19. Larsen, N. B. et al. Replication-Coupled DNA-Protein Crosslink Repair by SPRTN and the Proteasome in Xenopus Egg Extracts. Mol Cell 73, 574–588.e7 (2019).

20. Sparks, J. L. et al. The CMG Helicase Bypasses DNA-Protein Cross-Links to Facilitate Their Repair. Cell 176, 167–181.e21 (2019).

21. Gallina, I. et al. The ubiquitin ligase RFWD3 is required for translesion DNA synthesis. Mol Cell 81, 442–458.e9 (2021).

22. Borgermann, N. et al. SUMOylation promotes protective responses to DNA-protein crosslinks. EMBO J 38, e101496 (2019).

23. Leng, X. & Duxin, J. P. Targeting DNA-Protein Crosslinks via Post-Translational Modifications. Front Mol Biosci 9, 944775 (2022).

24. Stingele, J., Schwarz, M. S., Bloemeke, N., Wolf, P. G. & Jentsch, S. A DNA-dependent protease involved in DNA-protein crosslink repair. Cell 158, 327–338 (2014).

25. Ruggiano, A. & Ramadan, K. DNA-protein crosslink proteases in genome stability. Commun Biol 4, 11 (2021).

26. Stingele, J. et al. Mechanism and Regulation of DNA-Protein Crosslink Repair by the DNA-Dependent Metalloprotease SPRTN. Mol Cell 64, 688–703 (2016).

27. Vaz, B. et al. Metalloprotease SPRTN/DVC1 Orchestrates Replication-Coupled DNA-Protein Crosslink Repair. Mol Cell 64, 704–719 (2016).

28. Tsherniak, A. et al. Defining a Cancer Dependency Map. Cell 170, 564–576.e16 (2017).

29. Maskey, R. S. et al. Spartan deficiency causes genomic instability and progeroid phenotypes. Nat Commun 5, 5744 (2014).

30. Reinking, H. K., Hofmann, K. & Stingele, J. Function and evolution of the DNA-protein crosslink proteases Wss1 and SPRTN. DNA Repair (Amst) 88, 102822 (2020).

31. Li, F., Raczynska, J. E., Chen, Z. & Yu, H. Structural Insight into DNA-Dependent Activation of Human Metalloprotease Spartan. Cell Rep 26, 3336–3346.e4 (2019).

32. Reinking, H. K. et al. DNA Structure-Specific Cleavage of DNA-Protein Crosslinks by the SPRTN Protease. Mol Cell 80, 102–113.e6 (2020).

33. Centore, R. C., Yazinski, S. A., Tse, A. & Zou, L. Spartan/C1orf124, a reader of PCNA ubiquitylation and a regulator of UV-induced DNA damage response. Mol Cell 46, 625– 635 (2012).

34. Davis, E. J. et al. DVC1 (C1orf124) recruits the p97 protein segregase to sites of DNA damage. Nat Struct Mol Biol 19, 1093–1100 (2012).

35. Mosbech, A. et al. DVC1 (C1orf124) is a DNA damage-targeting p97 adaptor that promotes ubiquitin-dependent responses to replication blocks. Nat Struct Mol Biol 19, 1084–1092 (2012).

36. Kim, M. S. et al. Regulation of error-prone translesion synthesis by Spartan/C1orf124. Nucleic Acids Res 41, 1661–1668 (2013).

37. Zhao, S. et al. A ubiquitin switch controls autocatalytic inactivation of the DNA-protein crosslink repair protease SPRTN. Nucleic Acids Res 49, 902–915 (2021).

38. Lessel, D. et al. Mutations in SPRTN cause early onset hepatocellular carcinoma, genomic instability and progeroid features. Nat Genet 46, 1239–1244 (2014).

39. Ruggiano, A. et al. The protease SPRTN and SUMOylation coordinate DNA-protein crosslink repair to prevent genome instability. Cell Rep 37, 110080 (2021).

40. Maskey, R. S. et al. Spartan deficiency causes accumulation of Topoisomerase 1 cleavage complexes and tumorigenesis. Nucleic Acids Res 45, 4564–4576 (2017).

41. Husnjak, K. & Dikic, I. Ubiquitin-binding proteins: decoders of ubiquitin-mediated cellular functions. Annu Rev Biochem 81, 291–322 (2012).

42. Kolla, S., Ye, M., Mark, K. G. & Rapé, M. Assembly and function of branched ubiquitin chains. Trends Biochem Sci 47, 759–771 (2022).

43. Mirdita, M. et al. ColabFold: making protein folding accessible to all. Nat Methods 19, 679– 682 (2022).

44. Yaneva, D. et al. The FANCJ helicase unfolds DNA-protein crosslinks to promote their repair. Mol Cell 83, 43–56.e10 (2023).

45. Donsbach, M. et al. A non-proteolytic release mechanism for HMCES-DNA-protein crosslinks. The EMBO Journal **n/a**, e113360 (2023).

46. Renz, C. et al. Ubiquiton-An inducible, linkage-specific polyubiquitylation tool. Mol Cell 84, 386–400.e11 (2024).

47. Christman, J. K. 5-Azacytidine and 5-aza-2′-deoxycytidine as inhibitors of DNA methylation: mechanistic studies and their implications for cancer therapy. Oncogene 21, 5483–5495 (2002).

48. Weickert, P., Dürauer, S., Götz, M. J., Li, H.-Y. & Stingele, J. Electro-elution-based purification of covalent DNA-protein cross-links. Nat Protoc 19, 2891–2914 (2024).

49. Lopez-Mosqueda, J. et al. SPRTN is a mammalian DNA-binding metalloprotease that resolves DNA-protein crosslinks. eLife 5, e21491 (2016).

50. Kojima, Y. et al. FAM111A protects replication forks from protein obstacles via its trypsin-like domain. Nat Commun 11, 1318 (2020).

51. Dokshin, G. A. et al. GCNA Interacts with Spartan and Topoisomerase II to Regulate Genome Stability. Dev Cell 52, 53–68.e6 (2020).

52. Serbyn, N. et al. The Aspartic Protease Ddi1 Contributes to DNA-Protein Crosslink Repair in Yeast. Mol Cell 77, 1066–1079.e9 (2020).

53. Xu, P. et al. Quantitative proteomics reveals the function of unconventional ubiquitin chains in proteasomal degradation. Cell 137, 133–145 (2009).

54. Carvalho, A. F. et al. High-yield expression in Escherichia coli and purification of mouse ubiquitin-activating enzyme E1. Mol Biotechnol 51, 254–261 (2012).

55. Renz, C. et al. Ubc13-Mms2 cooperates with a family of RING E3 proteins in budding yeast membrane protein sorting. J Cell Sci 133, jcs244566 (2020).

56. Hughes, C. S. et al. Single-pot, solid-phase-enhanced sample preparation for proteomics experiments. Nat Protoc 14, 68–85 (2019).

57. Waterhouse, A. et al. SWISS-MODEL: homology modelling of protein structures and complexes. Nucleic Acids Res 46, W296–W303 (2018).

58. Mitchell, A. L. et al. MGnify: the microbiome analysis resource in 2020. Nucleic Acids Res 48, D570–D578 (2020).

59. Eastman, P. et al. OpenMM 7: Rapid development of high performance algorithms for molecular dynamics. PLoS Comput Biol 13, e1005659 (2017).

60. Jumper, J. et al. Highly accurate protein structure prediction with AlphaFold. Nature 596, 583–589 (2021).

61. Gordon, J. C. et al. H++: a server for estimating pKas and adding missing hydrogens to macromolecules. Nucleic Acids Res 33, W368–371 (2005).

62. Case, D. A. et al. AmberTools. J. Chem. Inf. Model. 63, 6183–6191 (2023).

63. Tian, C. et al. ff19SB: Amino-Acid-Specific Protein Backbone Parameters Trained against Quantum Mechanics Energy Surfaces in Solution. J Chem Theory Comput 16, 528–552 (2020).

64. Izadi, S., Anandakrishnan, R. & Onufriev, A. V. Building Water Models: A Different Approach. J Phys Chem Lett 5, 3863–3871 (2014).

65. Mamatkulov, S. & Schwierz, N. Force fields for monovalent and divalent metal cations in TIP3P water based on thermodynamic and kinetic properties. J Chem Phys 148, 074504 (2018).

66. Li, P., Roberts, B. P., Chakravorty, D. K. & Merz, K. M. Rational Design of Particle Mesh Ewald Compatible Lennard-Jones Parameters for +2 Metal Cations in Explicit Solvent. J Chem Theory Comput 9, 2733–2748 (2013).

67. Van Der Spoel, D. et al. GROMACS: fast, flexible, and free. J Comput Chem 26, 1701– 1718 (2005).

68. Miller, B. R. I. et al. MMPBSA.py: An Efficient Program for End-State Free Energy Calculations. J. Chem. Theory Comput. 8, 3314–3321 (2012).

69. Valdés-Tresanco, M. S., Valdés-Tresanco, M. E., Valiente, P. A. & Moreno, E. gmx_MMPBSA: A New Tool to Perform End-State Free Energy Calculations with GROMACS. J. Chem. Theory Comput. 17, 6281–6291 (2021).

70. Nguyen, H., Roe, D. R. & Simmerling, C. Improved Generalized Born Solvent Model Parameters for Protein Simulations. J. Chem. Theory Comput. 9, 2020–2034 (2013).

71. Delaglio, F. et al. NMRPipe: a multidimensional spectral processing system based on UNIX pipes. J Biomol NMR 6, 277–293 (1995).

72. Lee, W., Tonelli, M. & Markley, J. L. NMRFAM-SPARKY: enhanced software for biomolecular NMR spectroscopy. Bioinformatics 31, 1325–1327 (2015).

73. Mulder, F. A., Schipper, D., Bott, R. & Boelens, R. Altered flexibility in the substrate-binding site of related native and engineered high-alkaline Bacillus subtilisins. J Mol Biol 292, 111– 123 (1999).

74. Hodgins, R., Gwozd, C., Arnason, T., Cummings, M. & Ellison, M. J. The tail of a ubiquitin-conjugating enzyme redirects multi-ubiquitin chain synthesis from the lysine 48-linked configuration to a novel nonlysine-linked form. J Biol Chem 271, 28766–28771 (1996).

75. Parker, J. L. & Ulrich, H. D. Mechanistic analysis of PCNA poly-ubiquitylation by the ubiquitin protein ligases Rad18 and Rad5. EMBO J 28, 3657–3666 (2009).

